# Spatial tumour-immune ecosystems shape the efficacy of anti-PD1 immunotherapy in primary cutaneous melanoma

**DOI:** 10.1101/2025.07.07.661465

**Authors:** Félix Pham, Marion Dufeu, Valentin Benboubker, Maxime Grimont, Amélie Lhorisson, Justine Berthet, Marie Donzel, Raphaël Schneider, Laurie Tonon, Anne-Claire Doffin, Félix Boivin, Simon Durand, Bertrand Dubois, Jonathan Lopez, Christophe Caux, Jenny Valladeau-Guilemond, Anaïs Eberhardt, Stéphane Dalle, Julie Caramel

## Abstract

Intra-tumoral heterogeneity in melanoma arises from dynamic cancer cell plasticity and underlies various mechanisms of immune escape. Here, we combined high-plex immunofluorescence imaging with spatially resolved transcriptomics to map the architecture of melanoma cell states and their interactions with the immune microenvironment in primary cutaneous tumours prior to adjuvant anti-PD1 immune checkpoint inhibitor (ICI) treatment. Computational analyses showed that melanoma cells organise into spatially restricted patches, with a preferential organisation of undifferentiated cells associated with poor ICI efficacy. Neighbouring immune cell composition varied according to cancer cell states, with a crucial involvement of specific subsets of tumour-associated macrophages, driven by signalling pathways involving tumour-derived and microenvironmental cues such as IFN-γ and hypoxia. Integrated spatial analyses further revealed tumour-immune ecosystems that stratify patient outcomes, delineating configurations either associated with ICI efficacy or metastatic relapse. These results uncover the spatial landscape of tumour ecosystems and identify signalling pathways as potential targets for improving the efficacy of ICI in melanoma.

**Highlights:**

- Melanoma cell states spatial organization is associated with aPD1 therapy efficacy
- Spatial organization of TAM subsets is a crucial determinant of ICI outcome
- Melanoma cells in different states co-localise with functionally distinct TAM subsets
- Identification of cell-cell communication pathways that underlie tumour-TAM crosstalk
- A balance between tumour-immune ecosystems is associated with aPD1 therapy efficacy

## INTRODUCTION

Immune checkpoint inhibitors (ICI) targeting PD-1 have significantly improved the outcome of patients with advanced melanoma, yet primary or acquired resistance remains a clinical challenge for over half of patients treated (Wolchok et al. 2025; Eggermont et al. 2020). T cells subsets are key mediators of response to ICI (Sade-Feldman et al. 2018; Gide et al. 2019; Siddiqui et al. 2019; Mahuron et al. 2025), and their activation through cross-presentation by dendritic cells (DCs) is critical (Salmon et al. 2016; Barry et al. 2018; Gobbini et al. 2025). The presence of tertiary lymphoid structures (TLS) in tumours has also been linked to improved clinical outcomes, possibly due to their role in facilitating local immune priming and cellular interactions (Cabrita et al. 2020; Meylan et al. 2022). Moreover, recent research has highlighted the central role of tumour-associated macrophages (TAM) in shaping responses to ICI. Depending on their phenotype and spatial context, TAM may suppress or enhance anti-tumoral immune responses, ultimately tipping the balance towards resistance or responsiveness (Mulder et al. 2021; Bill et al. 2023; Li et al. 2024). Importantly, the spatial organization of the tumour immune microenvironment (TIME) is increasingly recognized as pivotal for anti-PD1 (aPD1) ICI efficacy. Specific spatial niches composed of CD8^+^ T cells (J. H. Chen et al. 2024), cDC1 (Gobbini et al. 2025) and/or TAM (Antoranz et al. 2022; H. Chen et al. 2025) have recently been described in several tumour contexts, emphasizing the functional relevance of tissue architecture in the regulation of the immune response. Yet, most studies have focused on immune-restricted parameters.

Although genetic determinants of ICI resistance have been described (Spranger, Bao, et Gajewski 2015; Van Allen et al. 2015; Zaretsky et al. 2016; Lim et al. 2023; Schiantarelli et al. 2025), the majority of melanoma tumours lack resistance-conferring genetic variants (Shain A. Hunter et al. 2015). Instead, melanoma exploits cancer cell plasticity as a key mechanism of therapeutic resistance (Arozarena et Wellbrock 2019; Marine, Dawson, et Dawson 2020), especially upon targeted therapies (Richard et al. 2016; Rambow et al. 2018). Similar to the epithelial-mesenchymal transition (EMT) that occurs in epithelial cancers, melanoma cell plasticity relates to the reversible phenotypic transitions between differentiated/melanocytic and undifferentiated/invasive cell states. These transitions, regulated by transcription factors, enable melanoma cell adaptation to their tumour microenvironment (Rambow, Marine, et Goding 2019), and drive intra-tumoral heterogeneity (ITH). Several immune escape mechanisms linked to melanoma cell plasticity have been described, suggesting a key contribution in shaping anti-tumoral immunity (Landsberg et al. 2012; Boshuizen et al. 2020; Benboubker 2022; Plaschka et al. 2022; Liu et al. 2024; Guetter et al. 2025). Nevertheless, the extent to which these mechanisms contribute to ICI resistance in patients remains largely unclear. While a single-cell RNA sequencing (scRNAseq) study recently deciphered the dynamic changes in melanoma cell states during aPD1 ICI in the metastatic setting (Pozniak et al. 2024), the spatial organization of melanoma cells, their interactions with the TIME and their clinical impact remain to be precisely addressed in primary cutaneous melanoma.

In this study, we investigated tumour-immune ecosystems in treatment-naïve primary cutaneous melanoma lesions from 57 stage III patients with lymph node metastasis having undergone surgery and adjuvant aPD1 ICI, by integrating RNA sequencing (RNA-seq), spatial multiplex immunofluorescence (mIF), and spatially resolved transcriptomics (SRT). Through the spatial analysis of melanoma ITH, we demonstrate that specific tumour-immune ecosystems emerge and are shaped by the crosstalk between melanoma cells and the immune microenvironment. Importantly, these spatially-defined ecosystems affect ICI efficacy and metastatic relapse. Our findings reveal a spatially-integrated tumour-immune architecture tuned by actionable interactions that contributes to therapeutic outcome in advanced melanoma.

## RESULTS

### Spatial organisation of differentiated and undifferentiated tumour cells within human primary cutaneous melanoma is associated with ICI efficacy

To investigate the heterogeneity and spatial organisation of melanoma cells according to their differentiation status in primary cutaneous melanoma lesions, we performed whole-slide mIF analysis, RNA-seq and SRT on tumour lesions from resectable stage III melanoma patients (n = 57) treated with wide local excision, complete lymph node excision when required, followed by adjuvant aPD1 ICI for 12 months with at least 48 months of follow-up from the diagnosis. The main clinical endpoints were the relapse status after the 2-year follow-up (relapse-free, RF or relapse, REL) and the recurrence-free survival (RFS) (Fig.1a). The demographic, clinical, histopathological and genetic information of each patient are presented in Table 1. Formalin-fixed, paraffin-embedded (FFPE) primary melanoma lesions analysed in this study were treatment-naïve, before any infusion of ICI. Importantly, stained sections were analysed as whole slides to comprehensively capture the heterogeneity of the lesions with single cell resolution.

**Figure 1:**
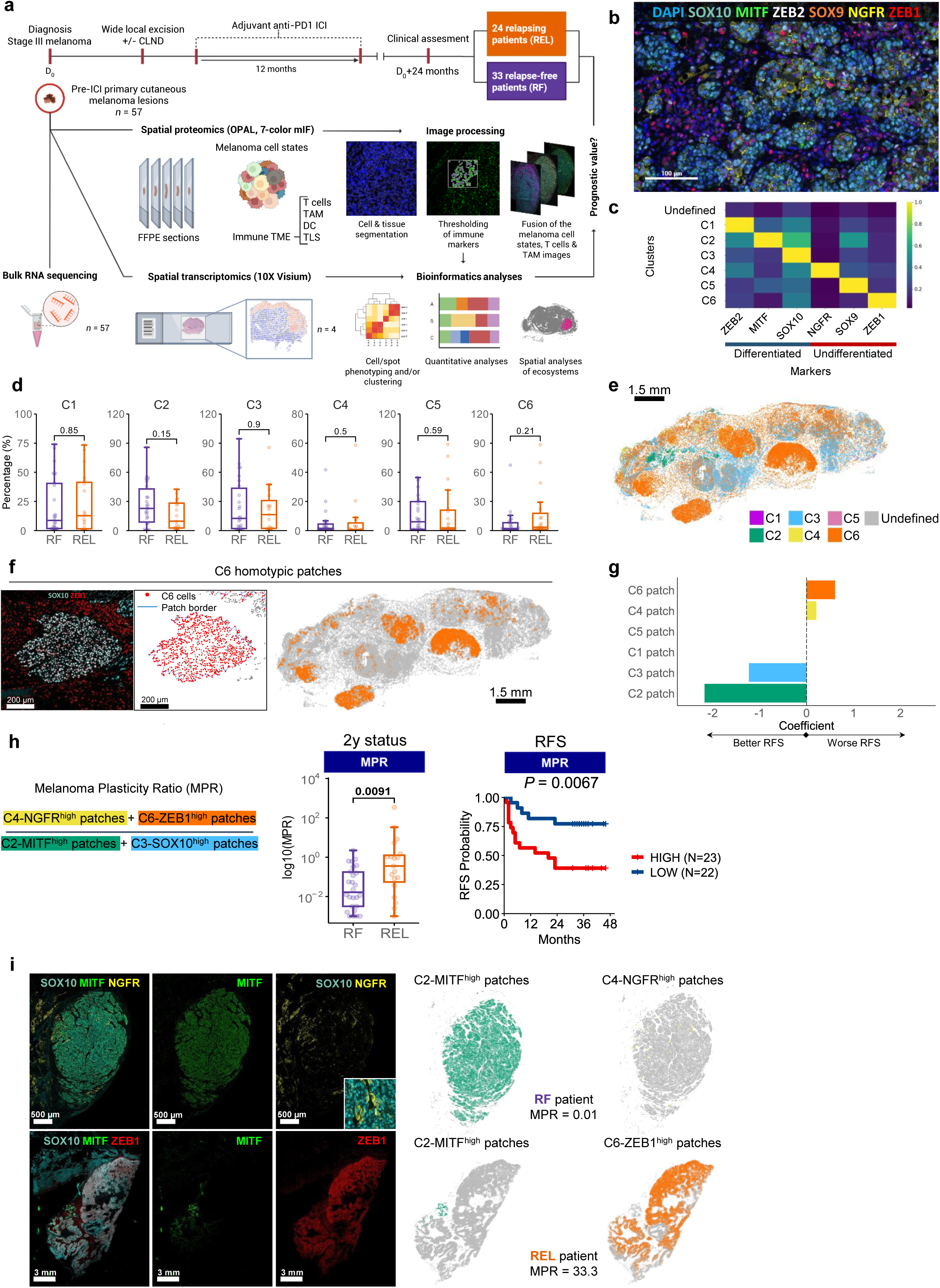
Spatial organization of differentiated and undifferentiated melanoma cells within primary tumours is associated with immune checkpoint inhibitor efficacy. **a** Schematic diagram of the analyses of pre-ICI stage III primary melanoma samples (*n* = 57 patients), showing the clinical follow-up above, and mIF, bulk-RNAseq and spatial transcriptomic analyses performed on formalin-fixed paraffin-embedded tissues (FFPE) below. CLND: complete lymph node dissection, DC: dendritic cells, ICI: immune checkpoint inhibitor, mIF: multiplex immunofluorescence, TAM: tumour-associated macrophages, TLS: tertiary lymphoid structure, TME: tumour microenvironment. Created with BioRender. **b** Representative tissue (of *n* = 45) section showing the mIF markers used to detect differentiated and undifferentiated melanoma cells. DAPI counterstaining appears as blue within each panel. Scale bar = 100 µM. **c** Unsupervised k-means clustering of each tumour cell (*n* = 7,644,292) from the sections stained in b to define 6 specific phenotypic clusters based on the mean fluorescent intensity (MFI) of each marker. **d** Boxplots showing the association between the proportion of each individual phenotypic cluster and ICI outcome at 2 years: relapse (REL, in orange) and relapse-free (RF, in purple). Mann-Whitney U tests, p-values are shown on the plot. **e** Spatial informatic reconstruction of the different phenotypic clusters in one tumor representative of n = 45. Scale bar = 1.5 mm. **f** mIF image of the C6 cells characterized by the co-expression of SOX10 and ZEB1 (left). These C6 melanoma cells (red) form homotypic patches (border in blue, middle) across the whole tumour (right). Left and middle scale bars = 200 µm. Right scale bar = 1.5 mm. **g** Coefficients of the features included in a Lasso regularized Cox proportional hazard model. Features with a positive coefficient are associated with worsened relapse-free survival (RFS), whereas those with a negative one are associated with improved RFS. **h** Definition of the Melanoma Plasticity Ratio (MPR) which is the ratio between undifferentiated C4+C6 cells on differentiated C2+C3 cells involved in a homotypic patch (left). Association between the MPR and the 2-y status (middle, boxplots, Mann-Whitney U tests) and RFS (right, Kaplan-Meier curves with MPR^high^ patients in red and MPR^low^ patients in blue, Log-rank test) of patients. **i** mIF image (left) and spatial reconstruction of the indicated clusters (right). First row: a mIF image of an RF-associated MPR^low^ tumour displaying few C4-NGFR^high^ patches (yellow) and numerous C2-MITF^high^ patches (green). Second row: a mIF image of a REL-associated MPR^high^ tumour displaying a high number of C6-ZEB1^high^ patches (orange) and only few C2-MITF^high^ patches (green).

**Table 1.** Clinical characteristics of the cohort.

| **2-year status** | **RF**<br>**N = 33 (57.9%)** | **REL**<br>**N = 24 (42.1%)** | **P value** | **Statistical test** |
| --- | --- | --- | --- | --- |
| **Age at diagnosis** |  |  | 0.81 | Wilcoxon |
| Years, mean +/- SD | 63.3 +/- 14.4 | 63.3 +/- 15.8 |  |  |
| **Follow-up duration** |  |  | 0.22 | Wilcoxon |
| Months, median (range) | 33 (27-42) | 32.5 (19-42) |  |  |
| **Sex** |  |  | 0.78 | Chi<sup>2</sup> |
| Female, n (%) | 11 | 10 |  |  |
| Male, n (%) | 23 | 16 |  |  |
| **BRAF/NRAS mutational status** |  |  | 0.64 | Chi<sup>2</sup> |
| BRAF, n (%) | 14 | 13 |  |  |
| NRAS, n (%) | 9 | 6 |  |  |
| Non-mutated<sup>§</sup>, n(%) | 10 | 5 |  |  |
| **Breslow index** |  |  | **0.02** | Wilcoxon |
| mm, median (range) | 3.4 (1.0-18.0) | 5.9 (1.2-20.0) |  |  |
| **Ulceration** |  |  | **0.03** | Chi<sup>2</sup> |
| Yes, n (%) | 16 (49) | 20 (83) |  |  |
| No, n (%) | 18 (51) | 5 (17) |  |  |
| **AJCC-8 stages** |  |  | **0.0047** | Chi<sup>2#</sup> |
| IIIA, n(%) | 5 (15) | 1 (4) |  |  |
| IIIB, n(%) | 9 (27) | 1 (4) |  |  |
| IIIC, n(%) | 17 (52) | 19 (79) |  |  |
| IIID, n(%) | 2 (6) | 3 (13) |  |  |
| **Location of metastases at relapse** |  |  | N/A | N/A |
| Skin | 0 | 6 |  |  |
| Lymph node | 0 | 12 |  |  |
| Lung | 0 | 11 |  |  |
| Liver | 0 | 7 |  |  |
| Surrenal glands | 0 | 4 |  |  |
| Spleen | 0 | 1 |  |  |
| Bones | 0 | 1 |  |  |
| Retroperitoneal | 0 | 1 |  |  |
| Brain | 0 | 3 |  |  |
| **Number of anti-PD1 infusions<sup>\*</sup>** |  |  | **<0.0001** | Wilcoxon |
| Median (range) | 17 (4-19) | 7 (2-18) |  |  |

<sup>§</sup>Or rare mutations: ARID2 (N = 1), KIT (N = 1), MAP3K1 (N=1), PTEN (N=1), RB1 (N=1), TERT (N=3)
<sup>\*</sup>Until first relapse
<sup>#</sup>Stages IIIA and IIIB were combined, and stages IIIC and IIID were combined for statistical purposes
AJCC: American Joint Commission on Cancer; N/A: not applicable; REL: relapse; RF: relapse-free;
SD: standard deviation

To explore the phenotypic heterogeneity of melanoma cells, a 7-color mIF panel composed of melanoma cell states markers was designed (Fig.1b and Extended Data Fig.1a). In addition to SOX10 (Cook et al. 2005; Capparelli et al. 2022), MITF (Carreira et al. 2006; Hoek et al. 2008; Rambow, Marine, et Goding 2019) and ZEB2 (Caramel et al. 2013; Vandamme et al. 2020) were selected as markers of a melanocytic/differentiated cell state. Conversely, ZEB1 (Caramel et al. 2013; Durand et al. 2024), NGFR (Tsoi et al. 2018; Boshuizen et al. 2020) and SOX9 (Tsoi et al. 2018; Wouters et al. 2020) were included as markers of invasive/neural crest stem-like (NCSC)/undifferentiated cell states.

Overall, 7,644,292 SOX10^+^ melanoma cells were detected across all tumour samples. Since these markers are not exclusive, we performed k-means clustering of melanoma cells based on the mean fluorescence intensity (MFI) of each marker. This analysis revealed six phenotypic clusters, that displayed a gradient in SOX10 expression with C3 exhibiting the highest SOX10 levels (Fig.1c and Extended Data Fig.1b). ZEB2 was expressed in C1 (MITF-int) and C2 (MITF-high) clusters, while NGFR expression was mostly restricted to C4 (Fig. 1c and Extended Data Fig.1b). A proportion of C6 (ZEB1^high^) and C5 (SOX9^high^) clusters displayed very low SOX10 levels (Fig. 1c and Extended Data Fig.1b), yet we were still able to detect these cells, thus identifying at least a subset of undifferentiated tumour cells that may have a downregulated SOX10 expression. Co-expression of SOX9 with MITF in C5 (Fig. 1c, Extended Data Fig.1b) may reflect a previously described intermediate cell state (Ennen et al. 2017; Wouters et al. 2020). A cluster presenting very low expression of all 6 markers was considered as undefined and removed from further analyses. For clarity, we hence referred to these key phenotypic clusters as C1-ZEB2^high^, C2-MITF^high^, C3-SOX10^high^, C4-NGFR^high^, C5-SOX9^high^ and C6-ZEB1^high^.

The proportion of each cluster was then quantified in all patients (Extended Data Fig.1c). Despite trends towards the enrichment in the baseline frequencies of the C2-MITF^high^ cluster in RF tumours and of the C6-ZEB1^high^ cluster in REL tumours, none of these phenotypic clusters was significantly associated with the 2-year relapse status (Fig.1d). This finding is similar to that of a recent study, where no baseline difference in melanoma cell states frequency was found between aPD1 ICI-responding and non-responding stage IV melanoma patients by scRNASeq (Pozniak et al. 2024). Remarkably, when considering the spatial organization of these tumours, melanoma cells belonging to the same phenotypic cluster tended to be spatially aggregated together (Fig.1e and Extended Data Fig.2a-i). To quantify this spatial organization, we defined homotypic tumour patches (Windhager et al. 2023) as regions containing at least five melanoma cells from the same phenotypic cluster within a 25 µm radius (Fig.1f). Interestingly, melanoma cells of the C2-MITF^high^ cluster predominantly formed homotypic patches, whereas cells with the C6-ZEB1^high^ phenotype were mostly isolated, which may reflect their invasive capacity (Extended Data Fig.3a).

Although the overall density of melanoma patches was not correlated with patient outcome after aPD1 treatment, a Cox regression model fitted with lasso regularization identified the most relevant cluster patches associated with RFS (Fig.1g and Extended Data Fig.3b-c). Notably, melanoma cells involved in C2-MITF^high^ and C3-SOX10^high^ patches were associated with better outcomes, while cells involved in C4-NGFR^high^ and C6-ZEB1^high^ patches with worse outcomes.

To further investigate the clinical relevance of this melanoma cell states organization, we calculated a melanoma plasticity ratio (MPR), defined as the ratio of C4-NGFR^high^ and C6-ZEB1^high^ melanoma cells organized as patches to C2-MITF^high^ and C3-SOX10^high^ cells in patches (Fig.1h). A higher MPR was found in REL patients and was consistently associated with worse RFS (Fig.1h-k). In contrast, a non-spatial MPR ratio that did not take into account tumour cell spatial organization was not significantly associated with ICI outcomes (Extended Data Fig.3d), highlighting the role of pre-treatment undifferentiated melanoma cells spatial distribution in reducing the efficacy of adjuvant aPD1 ICI. Interestingly, whereas the Breslow index –the most relevant histopathological prognostic metric– was significantly increased in REL patients (Table I), the prognostic value of the MPR ratio was independent of it (Extended Data Fig.3e-g).

To further quantify the diversity in melanoma cell clusters, we calculated the Shannon diversity index across every tumour, acknowledging that a higher index reflects a more diverse representation of every melanoma cell cluster (Spellerberg et Fedor 2003). Highly undifferentiated tumours exhibited a higher Shannon diversity index, suggesting that the acquisition of a more diverse melanoma cell state profile endows tumours with a survival advantage (Extended Data Fig.3h).

These findings revealed that melanoma cells are spatially organized in homotypic patches and that the ratio between undifferentiated and differentiated melanoma patches in the pre-treatment primary tumour influences the efficacy of aPD1 ICI.

### Integrative analysis of immune contexture highlights spatial organization of TAM subsets as a crucial determinant of ICI efficacy

We then performed a comprehensive characterization of the immune contexture using mIF across multiple tumour sections. This included assessing the densities and spatial organization of T cells (Extended Data Fig.4a, analysed with the CD3, CD8, PD1 and KI67 markers), DCs subpopulations (Extended Data Fig.4b) –conventional type 1 DCs (cDC1, CLEC9A^+^), conventional type 2 DCs (cDC2, CLEC10A^+^), plasmacytoid DCs (pDCs, BDCA2^+^), and migratory DCs (LAMP3^+^)–, TLS (Extended Data Fig.4c, assessed by the presence of at least one segregated T cell and B cell zones across the whole slide using the CD3, CD20 and DC-LAMP markers), and TAM (Fig.2a and Extended Data Fig.4d, analyzed with the pan-TAM marker CD68, pSTAT1 as a readout of proinflammatory TAM, CD163 as a marker of pro-tumoral TAM and PD-L1 expression).

**Figure 2:**
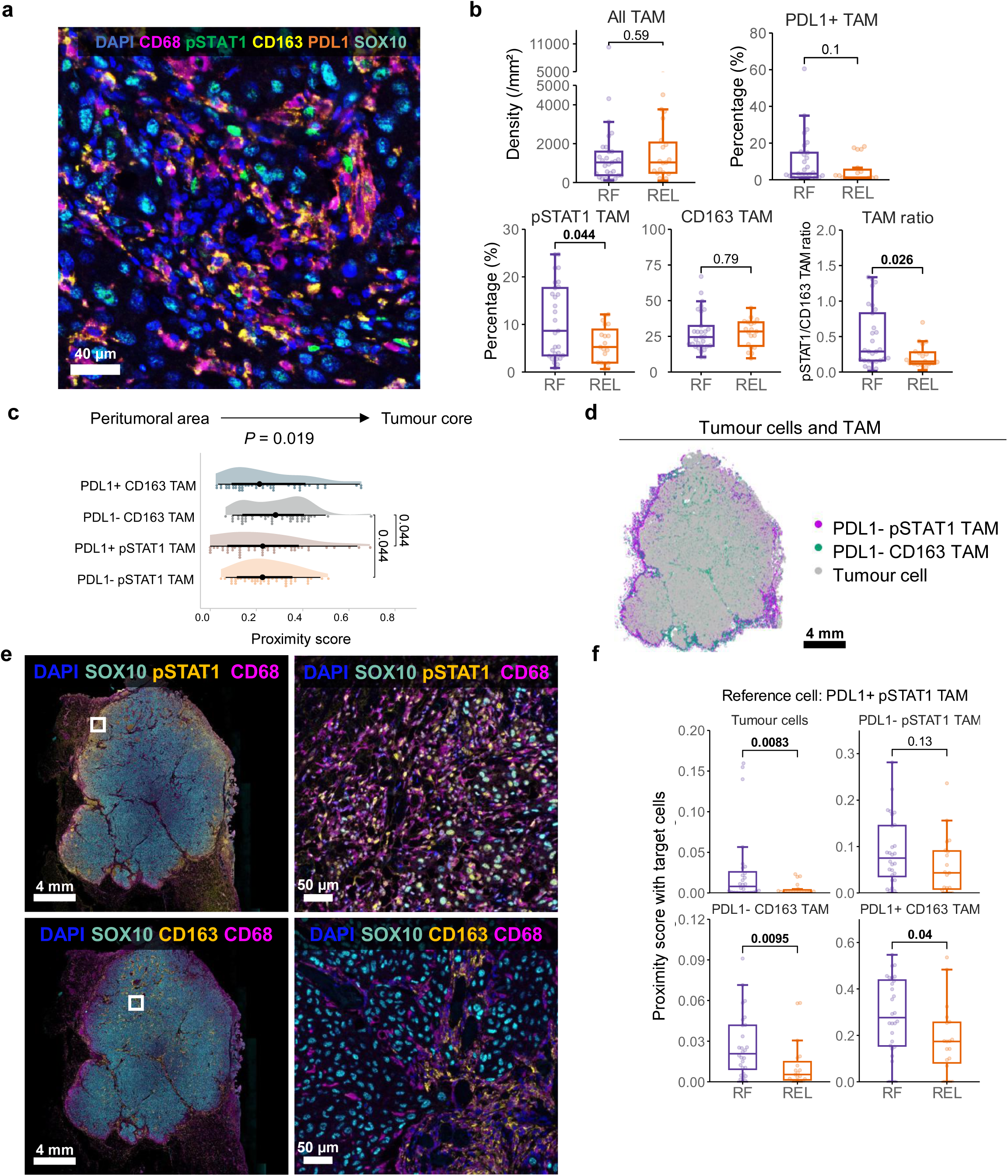
Tumour-associated macrophage density and spatial location dictate immune checkpoint inhibitors efficacy. **a** mIF antibodies used to study TAM subsets. Scale bar = 40 µM. **b** Density and percentage of TAMs in all tissue sections (*n* = 48). Graphs represent all TAM, PDL1^+^, pSTAT1^+^CD163^-^, and CD163^+^pSTAT1^-^ TAM, as well as the TAM ratio according to the 2-y status. Mann-Whitney U tests, p-values are indicated on the boxplots. **c** Proximity score illustrating the intra-tumoral infiltration by TAM subsets in a perimeter of 25 µm around a tumor cell. A higher score indicates more colocalization between both cell types independently of their frequencies. Friedman test and paired Wilcoxon tests, with p-values adjusted by Benjamini-Hochberg method. **d** Reconstructed tissue section displaying the specific spatial pattern of PDL1^-^pSTAT1^+^ TAM and PDL1^-^ CD163^+^ TAM subsets within the same tumour. Scale bar = 4 mm. **e** Corresponding mIF images from the tumour reconstructed in **(d)**. Right panels show enlarged views of inserts (scale bar = 50 µm) in the images on the left (scale bar = 4 mm). **f** Prognostic value of the proximity scores between PDL1^+^pSTAT1^+^ TAM and target cells in a radius of 25 µm. Mann-Whitney U tests between RF (purple) and REL (orange) patients. P-values are indicated in the boxplot.

Within the T cell compartment, we observed a trend towards increased CD8^+^ T cell density in RF patients, whereas REL patients exhibited a higher proportion of PD1^+^CD8^+^ T cells (Extended Data Fig.5a). We found that higher densities of cDC1, pDCs, and LAMP3^+^ DCs were associated with improved aPD1 efficacy (Extended Data Fig.5b), as we previously reported in stage IV melanoma (Gobbini et al. 2025). TLS were detected in only four of the tumours of our primary melanoma cohort, precluding any conclusion regarding their prognostic role. While total TAM (CD68^+^ cells) densities were comparable between RF and REL patients, we observed an increased proportion of pSTAT1^+^CD163^-^ TAM (pSTAT1 TAM) in RF patients (Fig.2b and Extended Data Fig.5c-d). Although no significant difference was detected in the relative proportion of CD163^+^pSTAT1^-^ TAM (CD163 TAM), the pSTAT1/CD163 TAM ratio was increased in RF patients, suggesting that pro-inflammatory TAM are associated with improved aPD1 efficacy (Fig.2b).

By integrating immune cell densities, we identified three distinct immunotypes at the patient level (Extended Data Fig.5e-f). Immunotypes 1 and 3 corresponded to previously described inflamed ("hot") and non-inflamed ("cold") TIMEs, respectively (Galon et Bruni 2019). More precisely, immunotype 1 was characterized by a high intra-tumoral infiltration of CD8^+^ and CD4^+^ T cells with DCs along with numerous pSTAT1 TAM and PDL1^+^ TAM. As expected, this density pattern was fuelled by a high IFN-γ signalling as evidenced by Gene Set Enrichment Analysis (GSEA) of RNA-seq data (Extended Data Fig.5e) (Versluis et al. 2024), with the upregulated genes from immunotype 1 (versus immunotype 3) being associated with immune cell recruitment and activation pathways (Extended Data Fig.5g-h). Immunotype 2 may represent an intermediate immune profile, as proposed in the ‘intermediate immunoscore’ model (Galon et Bruni 2019). Quantitative analyses of immune cell densities alone did not allow to accurately stratify patients according to their ICI outcome status (Extended Data Fig.5i), prompting us to further characterize the spatial organization of immune cells.

Accordingly, we performed spatial analyses using the normalized mixing score that quantifies the proximity between two given cell types, normalised by their respective proportion (Feng et al. 2023). These analyses revealed that PD1^+^CD8^+^ T cells were predominantly localized within the tumour core compared to other T cell subsets (Extended Data Fig.6a), and that their increased infiltration into the tumour correlated with increased aPD1 ICI efficacy (Extended Data Fig.6b). DCs tended to spatially cluster into homo- and heterotypic aggregates (Extended Data Fig.6c-e), with cDC1 cells being closer to the tumour core compared to other DC subsets (Extended Data Fig.6f). Comparisons between RF and REL patients demonstrated that the proximity of LAMP3^+^ DCs to CD8^+^ T cells or cDC2 cells was significantly correlated with a favourable prognosis (Extended Data Fig.6g).

The description of TAM heterogeneity and its association with ICI efficacy has emerged as a major focus of recent research but their spatial distribution still requires further analyses. Paired analyses within each tumour revealed that CD163 TAM and pSTAT1 TAM were spatially aggregated in different location, with PDL1^-^CD163 TAM being primarily located within the tumour core, whereas pSTAT1 TAM and PDL1^+^ TAM were more frequently found at the tumour margin (Fig.2c-e). Interestingly, an increased tumour infiltration by PDL1^+^pSTAT1 TAM was associated with favourable ICI outcomes, and their colocalization with CD163 TAM was similarly associated with enhanced aPD1 efficacy (Fig.2f). No other spatial interaction was significantly different between RF and REL patients.

Altogether, these integrated analyses of the immune landscape highlight novel patterns of immune cells spatial organization and raise important questions about how the spatial interactions between these immune cells and melanoma cell subsets affect the efficacy of aPD1 ICI.

### Differentiated and undifferentiated melanoma cells co-localize with functionally distinct macrophage populations

Given the highly organized spatial distribution of melanoma cells and immune contexture within the tumour, we hypothesized that the composition of the TIME may differ within these melanoma cell state-defined areas. To test this hypothesis, we integrated data from the three mIF panels —melanoma cell states, T cells and TAM— across 8 tumours containing concomitantly at least one large patch of differentiated (C2-MITF^high^ and/or C3-SOX10^high^) and undifferentiated (C4-NGFR^high^ and/or C6-ZEB1^high^) areas (Fig.3a-b). Differentiated and undifferentiated tumour cells related areas were manually contoured on the fused images and differential MPR ratio was confirmed before quantifying TAM and T cell densities within each area (Fig.3b-c). T cell densities did not significantly differ between differentiated and undifferentiated areas (Extended Data Fig.7a). However, TAM polarization exhibited a consistent shift, with a reduction in the CD163 TAM density and a concomitant increase in the pSTAT1 TAM density in differentiated areas as compared to the undifferentiated areas within the same tumour, leading to an elevated pSTAT1/CD163 TAM ratio in differentiated areas (Fig.3c and Extended Data Fig.7b).

**Figure 3:**
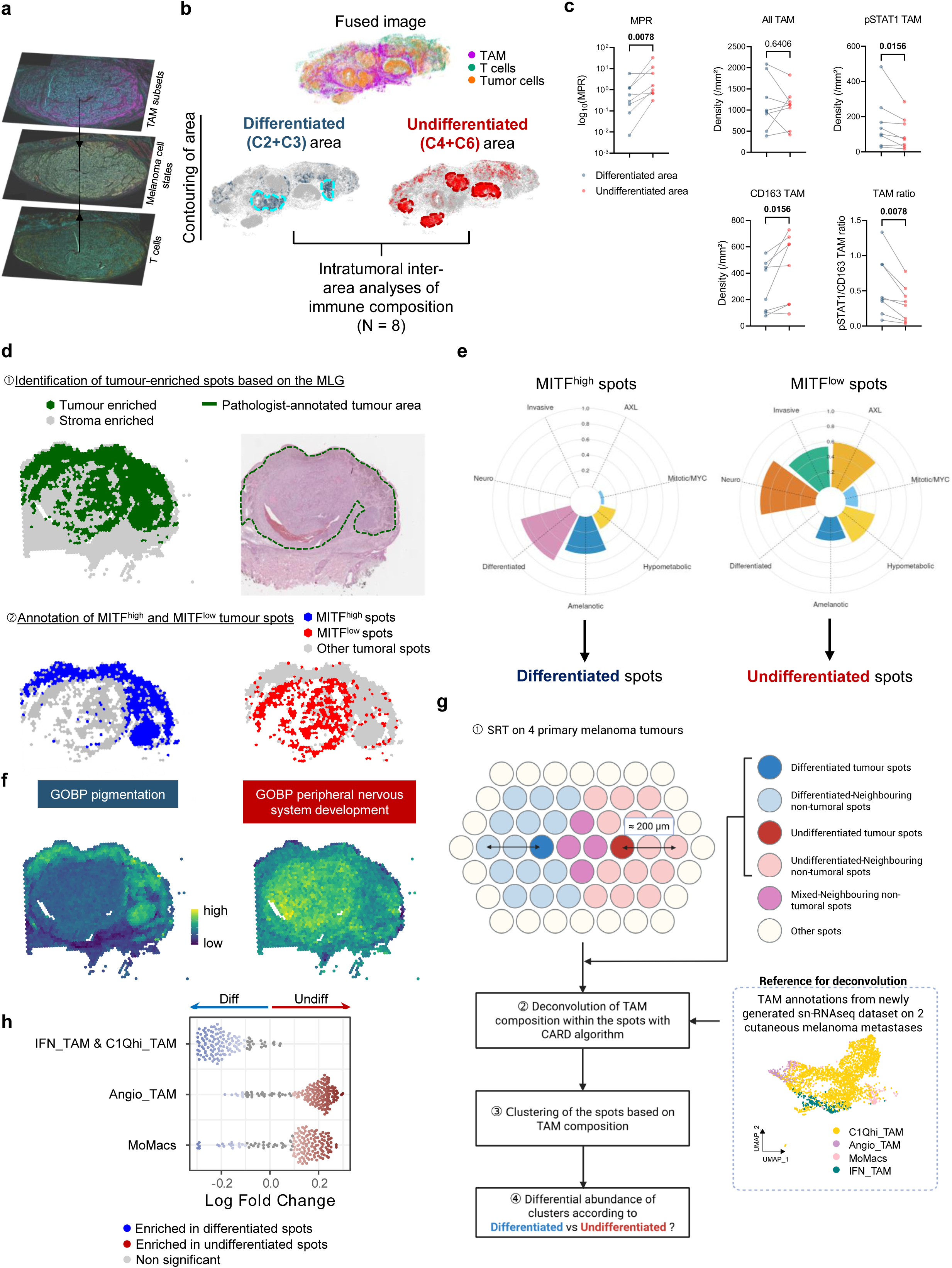
Melanoma cell states colocalize with specific TAM subsets. **a** Fused mIF images of *n* = 8 tumors showing TAM, T cells and melanoma cells, fused on a single image using HALO AI software. **b** Differentiated and undifferentiated tumour areas within the same tumour were manually contoured and the densities of T cell and TAM subsets within each area were calculated. **c** Paired comparisons of TAM subsets between differentiated and undifferentiated areas. Paired Wilcoxon test. P-values are indicated in the graphs. **d** Identification of tumour-enriched spots. First row: tumour-enriched spots labeled using the Melanoma Lineage Gene (MLG) signature from Pozniak et al, 2024 (left, binarized, with tumour-enriched spots in green), as compared to the pathological annotation of the tumour area (right, in dashed green). Second row: identification of MITF^high^ (left, in blue) and MITF^low^ (right, in red) spots based on the expression of *MITF*. **e** Radar plots characterizing tumor-enriched MITF^high^ and MITF^low^ spots compared to previously published gene signatures. **f** Spatial plot of Gene Ontology Biological Process (GOBP) pigmentation (blue) and peripheral nervous system development (red) in tumour-enriched spots. **g** Schematic diagram of the selection process of the tumour-enriched differentiated and undifferentiated spots and their neighboring spots (2 adjacent rows corresponding to an approximate perimeter of 200 µm) in tumour samples (*n* = 4). Deconvolution of TAM subsets within spots using the myeloid annotations from an in-house single-nuclei RNA sequencing (snRNAseq) dataset from 2 treatment-naïve cutaneous metastases. Uniform Manifold Approximation and Projection (UMAP) showing the annotations of the myeloid compartment in this dataset. The selected spots were then clustered to test for a differential abundance between differentiated and undifferentiated tumour-enriched spots and their neighbouring spots. Created with BioRender. **h** Milo differential abundance of TAM-enriched clusters between differentiated and undifferentiated tumour-enriched spots and their neighbouring spots. FDR (false discovery rate) threshold set at 0.1, with correction for multiple comparisons. Grey dots represent non-significant neighbourhoods according to the FDR threshold.

To explore in more detail the TIME in differentiated versus undifferentiated areas, we performed SRT analyses with the Visium platform on 4 out of the 8 previously described samples. Tumour-enriched spots were identified based on the expression of a previously established melanoma lineage gene (MLG) signature (Pozniak et al. 2024) and compared to manual pathological annotation, and subsequently stratified into MITF^high^ and MITF^low^ spots according to normalized *MITF* expression (Fig.3d and Extended Data Fig.8a). Differential pathway analyses confirmed that MITF^high^ spots largely overlapped with known differentiated or melanocytic signatures (Hu et al. 2024), thus defining differentiated tumour spots (Fig.3e). In contrast, MITF^low^ spots exhibited gene expression profiles consistent with undifferentiated states, including the NCSC, invasive, and EMT signatures, thereby delineating the undifferentiated tumour spots (Fig.3e and Extended Data Fig.8b). Gene Ontology Biological Process (GOBP) enrichment analyses further supported these distinctions, with pigmentation-related pathway being enriched in differentiated tumour spots, and peripheral nervous system development and neuron projection guidance pathways characterizing undifferentiated tumour spots (Fig. 3f and Extended Data Fig.8c).

To more precisely delineate TAM transcriptomic profiles within the spatial context of differentiated versus undifferentiated tumour spots, we performed cellular deconvolution of the spots using CARD algorithm (Y. Ma et Zhou 2022) (Fig.3g and Extended Data Fig.8d-e), leveraging an in-house single-nucleus RNAseq (snRNAseq) dataset from two cutaneous melanoma metastases, which is finely annotated for the myeloid compartment (R.-Y. Ma, Black, et Qian 2022). In line with the increase in pSTAT1 TAM in differentiated areas quantified by mIF, analyses of TAM transcriptomic profile within tumour spots and their neighbouring spots showed that differentiated tumour spots were enriched in C1QH TAM and interferon-primed TAM (IFN-TAM), a pro-inflammatory TAM subset overlapping with pSTAT1 TAM (Fig. 3h and Extended Data Fig.8f-h). Conversely, undifferentiated tumour spots were primarily adjacent to pro-angiogenic TAM (angio-TAM) (R.-Y. Ma, Black, et Qian 2022), a subtype of pro-tumoral TAM that expresses hypoxia-related genes and promotes metastatic dissemination and EMT (Fig. 3h and Extended Data Fig.8f-h) (Consonni et al. 2021; Cheng et al. 2021; R.-Y. Ma, Black, et Qian 2022).

Thus, these findings suggested that TAM polarization differs locally according to melanoma cell states, suggesting a preferential and spatially regulated crosstalk between TAM and melanoma cells.

### Tumoral and microenvironmental cues govern TAM polarization

To investigate the molecular cues originating specifically from the tumour cells, we performed ligand-receptor interaction analyses in the SRT data with CellChat (Jin, Plikus, et Nie 2025) focusing on cell-cell communications inferred between tumour-enriched and adjacent macrophage-enriched spots (Fig.4a-b and Extended Data Fig.9a). Deconvolution analyses confirmed the enrichment in TAM in these spots compared to other stroma-enriched spots (Extended Data Fig.9b). Differential analyses led to a list of ligand-receptors interactions specifically originating from differentiated or undifferentiated tumour spots (Fig. 4b). We next analysed in the single-cell RNA-seq dataset from Pozniak et al. (Pozniak et al. 2024) the expression pattern of the ligands in the distinct melanoma cell clusters (Fig.4c), as well as the level and specificity of expression of the receptors in TAM versus other cells from the TME (Extended Data Fig.10a). Among the top interactions enriched in differentiated spots was IL16-CD4 (Fig.4b). *IL-16* was highly expressed in differentiated spots (Fig.4d) and predominantly expressed by the “interferon αß response” melanoma cell subset (Fig.4c). IL-16 is known to enhance anti-tumoral immune response by stimulating the production of pro-inflammatory cytokines (Lynch et al. 2003). Even though the IL-16 receptor, *CD4*, is not only expressed by T cells but also by a proportion of TAM (Fig.4e and Extended Fig.10a), IL-16 was recently shown to reshape TAM phenotypes by altering CD4^+^ T cell function in the TME (Wen et al. 2025). Importantly, decreased IL-16 production was associated with resistance to ICI in breast and lung cancer patients (Wen et al. 2025). IFNγ-induced T-cell attracting chemokines *CXCL9*, *10* and *11*, which signal through the CXCR3 receptor, were also enriched in differentiated spots, and specifically expressed by the “Antigen presentation” melanoma cell subset (Fig.4c-d). This is reminiscent of our previous results showing that ZEB1-dependant decrease in CXCL10 melanoma cell-specific expression was associated to immune escape (Plaschka et al. 2022). Similarly, the CCL5-CCR5 T cell attracting signalling was enriched in differentiated spots, with *CCL5* being specifically enriched in the “Antigen presentation” melanoma cell subset (Fig.4c-e). Importantly, tumour-cell produced *CCL5* and *CXCL9* co-expression was previously associated with prolonged survival and better response to ICI (Dangaj et al. 2019). Since CXCL9, 10 and 11 are also strongly produced by TAM upon exposure to T cell-derived IFNγ, an amplification loop may further sustain this anti-tumoral immune ecosystem. Consistent with this, differentiated tumour spots exhibited a high expression of pSTAT1 signature and interferon gamma response pathways (Extended Data Fig.11a), reminiscent of the strong IFN-γ signalling observed in Immunotype 1 tumours enriched in pSTAT1 TAM (Fig. 2a).

**Figure 4:**
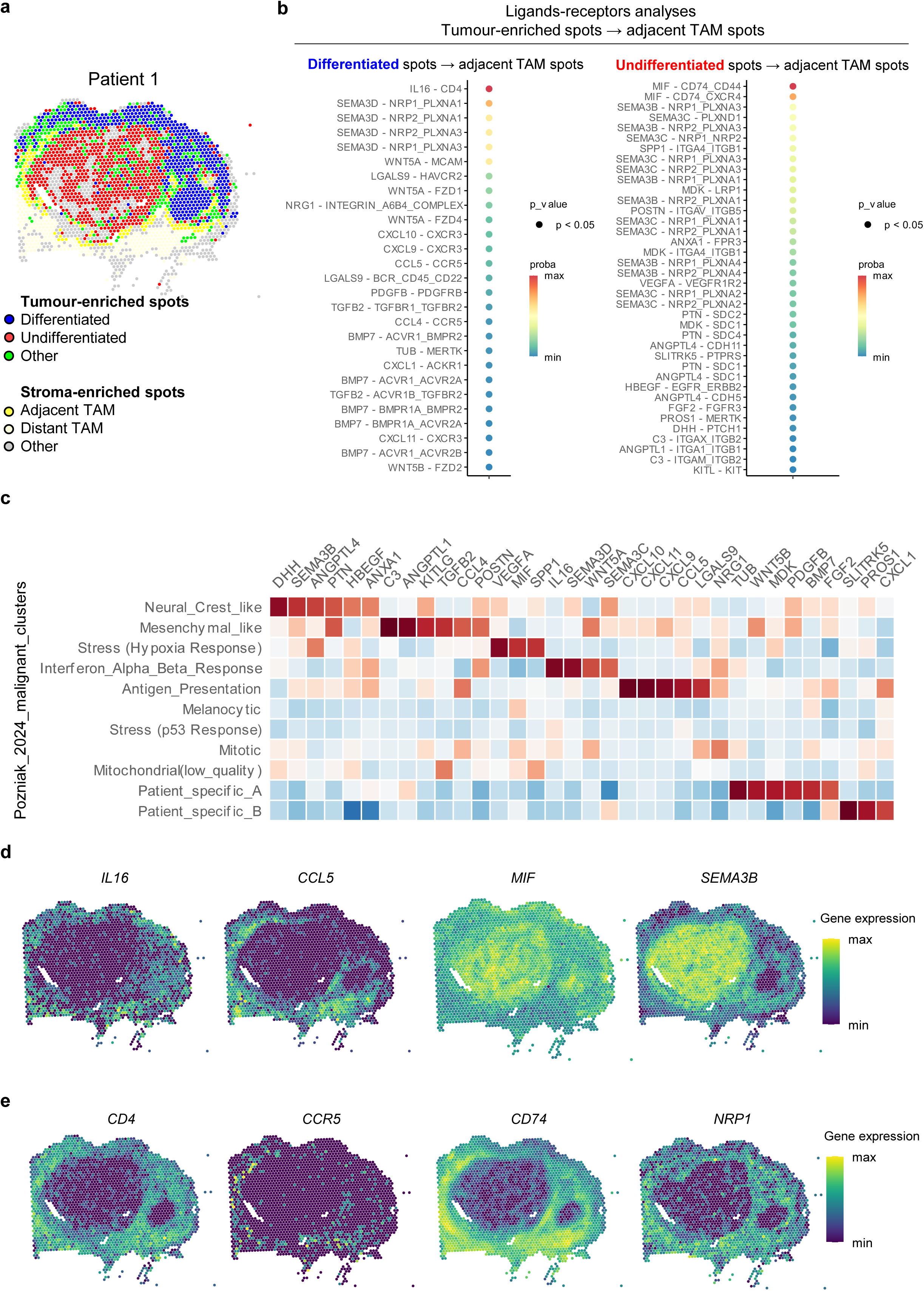
Differential cell-cell communication between melanoma cells and TAM according to the melanoma cell state. **a** Labeling of cell spots of patient #1, including adjacent TAM-enriched spots in yellow (TAM-enriched spots located in a radius of ≤ 200 µm to the nearest tumour-enriched spot) and distant TAM-enriched spots in light yellow (TAM-enriched spots located in a radius of > 200 µm to the nearest tumour-enriched spot). **b** Bubble plot diagrams illustrating the cell-cell communications, inferred by CellChat, between ligands from differentiated/undifferentiated tumour-enriched spots and receptors from adjacent TAM-enriched spots. Results from the 4 tumours. **c** Heatmap showing the gene expression as a z-score of the ligand-coding genes among malignant clusters (left) in the Pozniak et al (2024) dataset. **d** Spatialplot of ligand-coding genes *IL16* and *CCL5* (differentiated spots) and *SEMA3B* and *MIF* (undifferentiated spots) on patient #1 sample. **e** Spatialplot of receptor-coding genes *CD4* (receptor of IL16), *CCR5* (CCL5), *CD74* (MIF), and *NRP1* (SEMA3B) on patient #1 sample.

On the other side, within undifferentiated spots, various ligands already known to promote both melanoma cells invasive state and TAM polarization towards pro-tumoral states, as well as tumour angiogenesis and resistance to ICI (de Visser et Joyce 2023), were upregulated, with on top of the list, *MIF* (macrophage migration inhibitory factor) and its receptor *CD74* (Fig.4b-e). *MIF* is strongly expressed by “stress hypoxia” melanoma cells and the targeting of MIF-CD74 signaling on TAM was shown to restore anti-tumor immune response in melanoma (Figueiredo et al. 2018; de Azevedo et al. 2020). Semaphorins and their receptors Neuropilins and Plexins may also play a central role at the crossroad between TAM attraction and angiogenesis. The *SEMA3B* isoform was indeed specifically increased in the “NCSC” and “mesenchymal-like” melanoma cell subsets (Fig. 4c-d), while expression of its receptor *NRP1* by TAM (Fig.4e and Extended Data Fig.10a) was previously shown to regulate their entry into hypoxic niches (Casazza et al. 2013). Consistently, undifferentiated tumour spots displayed a hypoxic environment (Di Giovannantonio et al. 2025) (Extended Data Fig.11a), along with enrichment of the angiogenic VEGFA-VEGFR1R2 pathway (Inoue et al. 2002; Comunanza et al. 2023) (Extended Data Fig.11b-c). Notably, *VEGFA* is specifically expressed by “stress hypoxia” melanoma cells (Fig.4c). Other ligands enriched in “NCSC” and “mesenchymal-like” cells, the signaling of which was known to directly regulate TAM polarization, notably included Midkine (*MDK*) and is receptor *LRP1* (Cerezo-Wallis et al. 2020; Catena et al. 2025) (Extended Data Fig.11b-c).

Overall, integrating spatially resolved protein-based, transcriptomic and cell-cell communication analyses uncovered that distinct melanoma cells states differentially engage with TAM within spatially-conserved areas to modulate their polarization. Beyond TAM, these melanoma cell subsets likely also interact with other immune cells –particularly T cells– thereby potentially shaping functionally divergent tumour-immune ecosystems.

### Distinct immune ecosystems emerge around melanoma cell state patches

To validate that distinct tumour-immune ecosystems coexist within primary melanoma tumours, we integrated data from our three mIF panels, namely melanoma cell states, T cells and TAM, as described above (Fig.3a), but across the 45 tumours. To investigate the immediate surrounding of melanoma subsets, the borders of C2-MITF^high^, C3-SOX10^high^, C4-NGFR^high^ and C6-ZEB1^high^ patches were then computationally extended by 50 µm, defining an immune cellular neighbourhood (CN) (Fig.5a). Louvain clustering of 24,082 CNs based on their immune cell composition identified 7 specific and recurrent CNs (Fig.5b-d and Extended Data Fig.12a-b). Interestingly, differentiated (C2-MITF^high^, C3-SOX10^high^) and undifferentiated (C4-NGFR^high^ and C6-ZEB1^high^) patches displayed preferential immune CNs: while differentiated patches were enriched in CN3 and CN7 (Fig.5e) containing PDL1^+^ TAM co-occurring with CD8^+^PD1^+^ T cells, undifferentiated patches were enriched in CN5 characterized by the colocalization of PDL1^-^CD163^+^ TAM with CD8^+^PD1^-^ T cells, and in CN6 enriched in PDL1^-^ TAM (Fig.5e and Extended Data Fig.12c).

**Figure 5:**
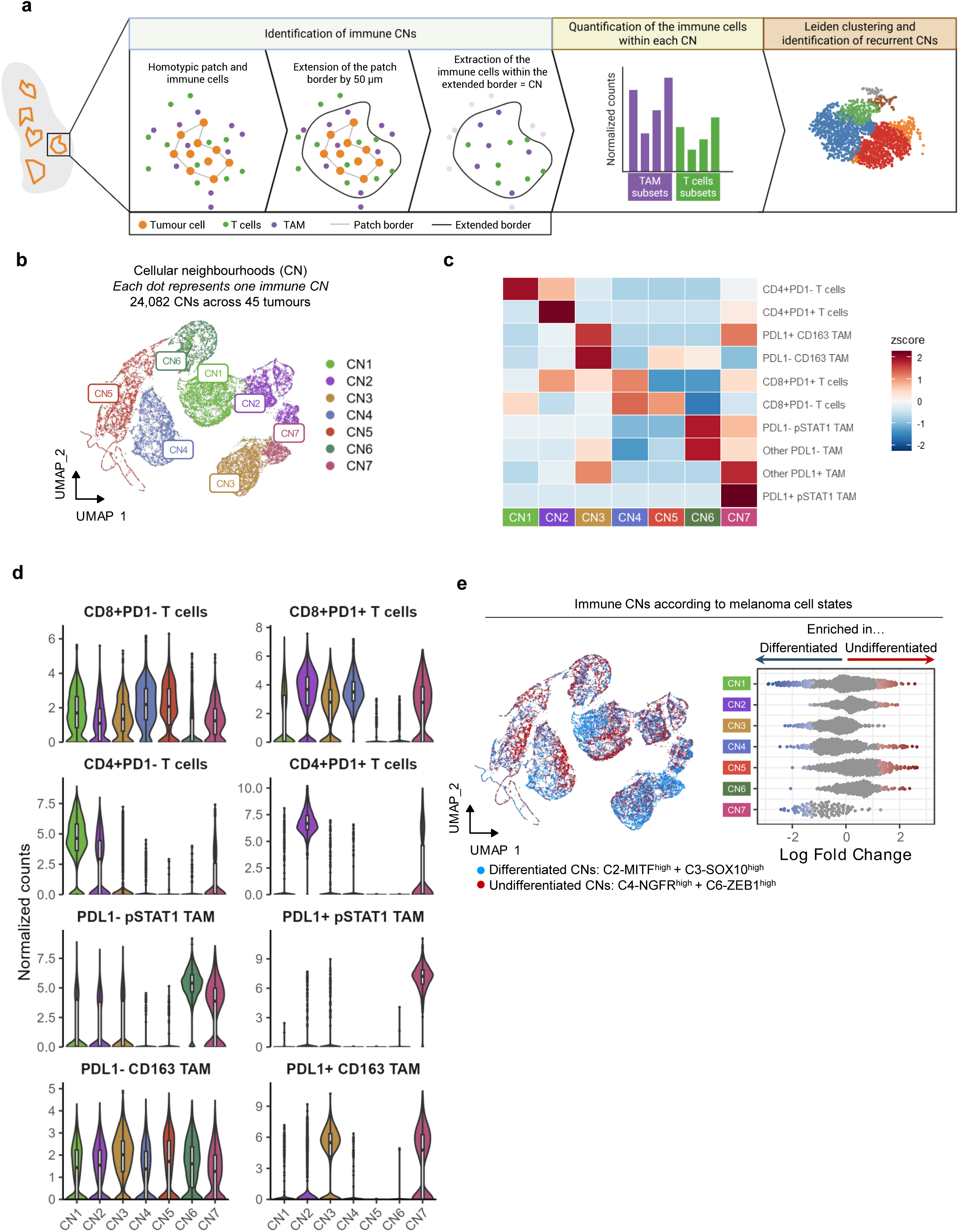
Crosstalk between melanoma cell subsets and immune cell neighbourhoods defines different ecosystems. **a** Schematic diagram of the analysis pipeline for the identification of immune cellular neighborhoods (CNs) around each C2/C3/C4/C6 tumor patch in fused mIF melanoma cells, T cells, and TAM images from 45 tumour samples. Created with BioRender. **b** UMAP of the Louvain clustering of the *n* = 24,082 CNs across the 45 samples. Each dot represents one CN. **c** Heatmap of the most variable cell types across the 7 identified CNs. Colours depend on the z-score of the most variable cell types between CNs. **d** Violin boxplots of the normalized count of T cell and TAM subsets in the different CNs. The counts were normalized against the area of each CN and against the proportion of the given cell type across the whole slide. **e** UMAP (left) and beeswarmplot (right) of the differential abundance of CNs between differentiated (C2/C3, in blue) and undifferentiated (C4/C6, in red) patches. FDR (false discovery rate) threshold set at 0.1, with correction for multiple comparisons.

Overall, our results suggest that the patches of distinct melanoma cell states are associated with specific immune neighbourhoods, hence defining preferred ecosystems.

### Spatial tumour-immune ecosystems dictate the efficacy of ICI in melanoma

We next investigated whether these ecosystems influence the efficacy of aPD1 ICI. Supporting this hypothesis, the seven CNs displayed differential enrichment between REL and RF patients, with CN3 and CN7 associated with favourable outcomes, whereas CN5 and CN6 were enriched in REL patients (Fig. 6a). While CN analyses highlight distinct microenvironmental ecosystems associated with patient outcomes, this approach was guided by predefined hypotheses centred on melanoma cell-centric neighbourhoods. To gain a broader and more comprehensive understanding of the spatial determinants of prognosis, we therefore complemented this analysis with an entropy-based framework to systematically address the prognostic impact of every possible pairwise colocalizations between melanoma cell states, T cells and TAM subsets (Fig. 6b). For this, we used a previously published analytical pipeline (Feng et al. 2023) that quantified the colocalization between these cell types, weighted for their respective density, to generate a unique colocalization score that accounts for inter-sample heterogeneity (See Methods and Fig. 6b). Through this approach, we identified specific pairwise colocalizations associated with either improved or worsened RFS following adjuvant aPD1 ICI (Extended Data Fig. 13a). Strikingly, the PDL1^+^pSTAT1 TAM was the cell type for which the colocalization score was the most frequently linked to better RFS. Interestingly, despite similar overall densities of CD163 TAM in RF and REL patients (Fig.2b), high colocalization between PDL1^-^CD163 TAM and either CD8^+^PD1^-^ T cells or C3-SOX10^high^ melanoma cells was significantly associated with a poorer survival, suggesting an immunosuppressive role played by this TAM subset.

**Figure 6:**
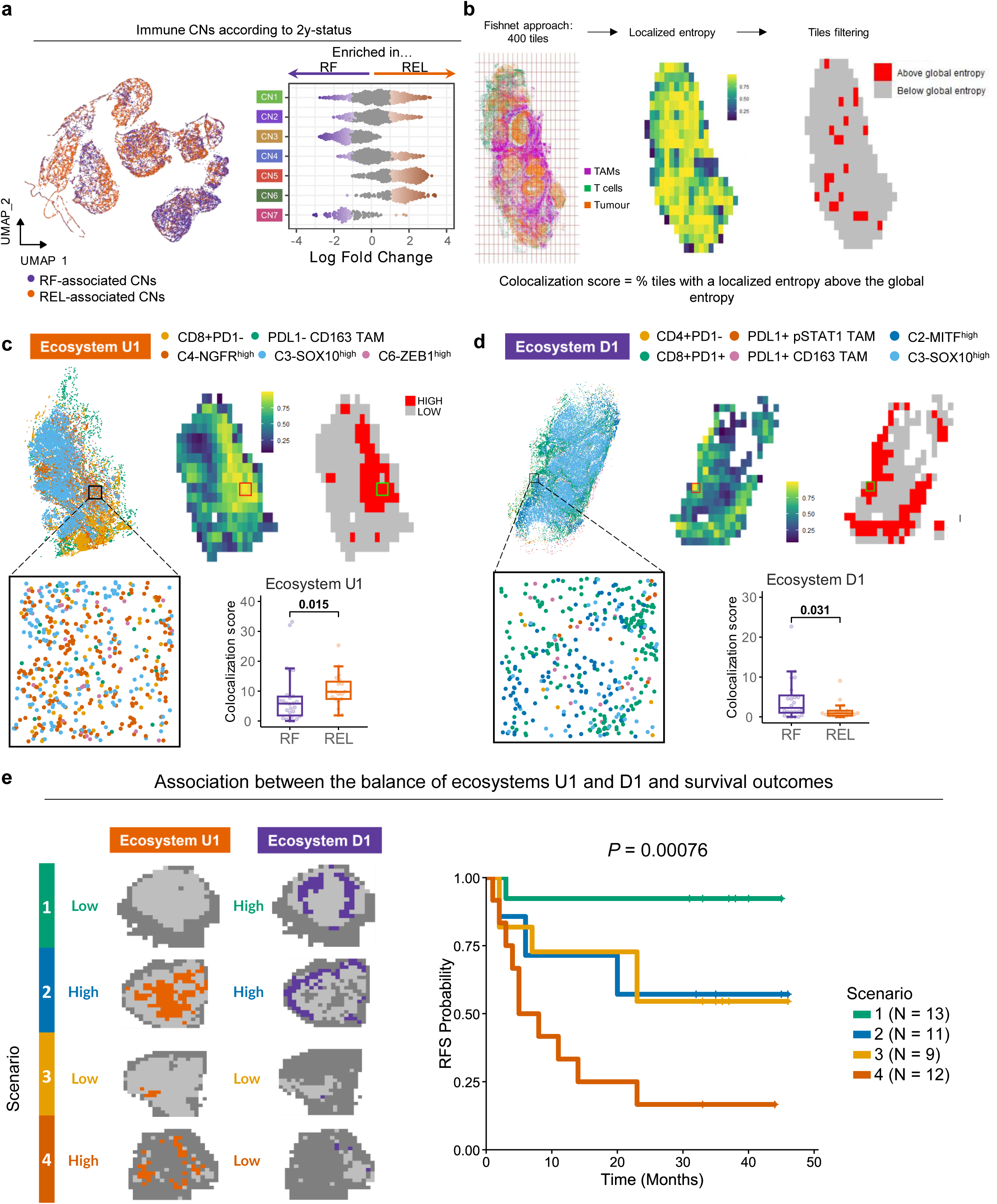
Tumour-immune spatial co-organization defines functional ecosystems driving ICI efficacy. **a** UMAP (left) and beeswarmplot (right) of the differential abundance of CNs between RF-associated (in purple) and REL-associated (in orange) patches. FDR (false discovery rate) threshold set at 0.1, with correction for multiple comparisons. **b** Schematic diagram of the pipeline used to calculate the co-localization score of given cell types. Fishnet approach to divide each tumor into a grid containing 20*20 tiles, excluding empty squares to calculate the localized entropies between TAM, T cells and melanoma subsets. As an example, localized entropies of a pattern of two cell types (TAM and tumour cells) are displayed and compared to the global entropy calculated on the whole tumour. The co-localization score is the percentage of tiles with a localized entropy above the global entropy for a given pattern. **c** Ecosystem U1 composition and prognostic impact (*n* = 45). In the upper row, the left panel represents one point per cell, the middle panel is the binned representation of the localized entropy of U1 cell types, and the right panel represents in red the tiles with a localized entropy above that of the global entropy of the given tumor (hotspots). In the lower row, an enlarged view of one area of high co-localization and boxplots of the colocalization score according to the 2-year status REL (relapse) or RF (relapse-free), of patients. Mann Whitney U test: the p-value is indicated on the graph. **d** Ecosystem D1 composition and its prognostic impact, as described in B for the U1 ecosystem. Mann-Whitney U test. **e** Description of the 4 main scenarios according to the proportion of U1 and D1 ecosystems across the 45 tumours (left) and their Kaplan Meier curves (right). Log-rank test, the p-value is indicated in the graph.

Given the frequent recurrence of specific pairwise colocalizations, we hypothesized that these interactions likely represented larger spatial ecosystems involving multiple heterotypic cell types. To explore this, we focused on cell types involved in detrimental colocalizations, i.e C6-ZEB1^high^, C4-NGFR^high^, C3-SOX10^high^ melanoma cells, mostly corresponding to undifferentiated tumour areas, CD8^+^PD1^-^ T cells and PDL1^-^CD163 TAM. Entropy-based colocalization analyses revealed the presence of this undifferentiated-1 (U1) ecosystem across 43 (out of 45) tumours, which was indeed enriched in REL-associated tumours (Fig. 6c). Conversely, through identification of recurrent pairwise interactions associated with favourable outcomes across the tumours, we identified the differentiated-1 (D1) ecosystem that was associated with a prolonged RFS, and was composed of C2-MITF^high^, C3-SOX10^high^, PDL1^+^pSTAT1, PDL1^+^CD163 TAM, CD8^+^PD1^+^ and CD4^+^PD1^-^ T cells (Fig. 6d and Extended Data Fig. 13b-c).

Most of the patients displayed both U1 and D1 ecosystems: thus, we hypothesized that the relative balance between these ecosystems could influence aPD1 efficacy. Indeed, we identified four distinct prognostic scenarios based on their frequency (Fig. 6e). The best-case scenario implied only few hotspots of the U1 ecosystem associated with a high prevalence of the D1 ecosystem (n = 13, n = 1 REL) whereas the worst-case scenario consisted of a high number of hotspots of the U1 ecosystem associated with only few to none of the D1 ecosystem (n = 12, n = 9 REL). In contrast, two opposite scenarios converged to the same intermediate prognosis: either the tumour was roughly equally enriched in U1 and D1 ecosystems (n = 11, n = 3 REL) or both ecosystems were scarcely present across the tumour (n = 7, n = 3 REL). The clinical relevance of these ecosystems was further highlighted by their ability to predict the efficacy of ICI with higher accuracy than the Breslow index, the AJCC-8 staging, the MPR alone or the immunotypes (Extended Data Fig. 13c). Interestingly, through regularized Cox regression on the MPR, the immunotypes, the ecosystems, and their interactions terms, we computed a composite score that yielded a high predictive value in discerning REL and RF patients, underscoring the clinical relevance of combining tumoral, immune and spatial parameters (Extended Data Fig. 13d-e).

Overall, these findings suggest that distinct tumour-immune ecosystems coexist in spatially specific areas of melanoma tumours, and that their balance is a key determinant of aPD1 ICI efficacy.

## DISCUSSION

In this study, we leveraged integrative RNA-seq, mIF, and SRT analyses from treatment-naïve stage III melanoma patients treated with ICI to explore the spatial organization of intra-tumoral heterogeneity and immune cell contexture, and its impact on patient outcome. Our findings revealed non-random spatially arranged, tumour-immune ecosystems driven by heterotypic tumour-immune crosstalks which correlate with aPD1 ICI efficacy.

Although prior studies have demonstrated the implication of melanoma cell plasticity in immune escape (Landsberg et al. 2012; Boshuizen et al. 2020; Benboubker 2022; Plaschka et al. 2022; Guetter et al. 2025), dedifferentiated melanoma cells found in metastases has only recently been reported to predict resistance to aPD1 ICI following a single treatment cycle (Pozniak et al. 2024). Whether such undifferentiated melanoma populations contribute to ICI efficacy in the pre-treatment setting, and in primary cutaneous melanoma has remained unclear. Here, whole-slide spatial profiling of treatment-naïve primary cutaneous melanomas revealed that melanoma cells sharing phenotypic identity co-localize together into homotypic spatial patches, mirroring the architecture observed in lymph node metastases (Pozniak et al. 2024). Notably, an enrichment in undifferentiated NGFR^high^ and ZEB1^high^ patches – relative to differentiated MITF^high^ and SOX10^high^ patches – prior to treatment was associated with a decreased ICI efficacy. Interestingly, the same ratio of melanoma cells without accounting for these homotypic patches was not predictive of treatment outcomes, highlighting the importance of spatial context. These findings suggest that the formation of homotypic tumour cell patches, rather than cellular composition alone, modulates therapeutic response, with the balance between distinct melanoma cell states serving as a potential determinant of ICI efficacy.

The non-random arrangement of melanoma cell states implies that signals originating from the TME – such as chemokines or cytokines, namely IFN signalling – orchestrate the emergence of these tumoral homotypic patches (Pozniak et Marine 2025). Conversely, the organization of these patches may, in turn, influence the phenotypic states of surrounding immune and stromal populations, suggesting a crosstalk within the TME that may critically shape therapeutic outcomes. This is illustrated through the unsupervised clustering of the immune cell composition surrounding these homotypic melanoma patches that unveiled recurrent immune CNs. Remarkably, differentiated melanoma patches were more frequently surrounded by CNs composed of PDL1^+^ TAM and PD1^+^CD8^+^ T cells, compared to undifferentiated melanoma patches. This finding is reminiscent of the colocalization of interferon-responsive tumour cell states with macrophages and T cells observed in a pan-cancer analysis (Barkley et al. 2022). Additionally, the intra-tumoral analyses of TAM phenotypes identified the colocalization between CD163 TAM and undifferentiated melanoma niches that mirrored observations in primary acral lentiginous melanoma where M2-like APOE^+^CD163^+^ TAM promote an undifferentiated melanoma cell state through the IGF1-IGF1R signalling pathway in spatially restricted regions (Liu et al. 2024). Here, cell-cell communication analyses performed on spatial transcriptomic data uncovered several pathways through which differentiated and undifferentiated melanoma cell subsets might shape the phenotype of neighbouring TAM. Notably, IL-16 and CCL5 produced by differentiated melanoma cells may promote proinflammatory TAM, through their dialogue with T cells, leading to a local amplification of IFNγ signalling. In contrast, in hypoxic regions enriched in undifferentiated melanoma cells, decreased production of IL-16/CCL5 by tumour cells and IFN-dependant chemokines by tumour cells and TAM, coincides with increased production of a range of pro-invasive and pro-angiogenic molecules such as MIF, SEMA3-B, MDK or VEGFA, that may altogether favour TAM polarisation towards pro-tumour subsets. In addition to this crosstalk, our spatial transcriptomic data highlighted how spatially constrained microenvironmental cues, such as a hypoxia gradient, might engage both the tumoral and non-tumoral compartments altogether.

Collectively, these findings support the growing paradigm of tumour ecosystems (X. Chen et Song 2022), whereby heterotypic interactions between melanoma cell states and immune populations modulate the efficacy of ICI. First, entropy-based analyses identified PDL1^+^ pSTAT1 TAM as the cell population most consistently associated with favourable ICI outcomes. This result goes beyond prior reports implicating pro-inflammatory, “M1-like” TAM in enhancing ICI response within spatially restricted tumour regions (Antoranz et al. 2022; J. H. Chen et al. 2024; H. Chen et al. 2025). Indeed, the D1 ecosystem, enriched in RF patients, and marked by the previously reported colocalization of CD8^+^PD1^+^ T cells with PDL1^+^ pSTAT1 TAM (Antoranz et al. 2022), was found in preferential interactions with differentiated MITF^high^ and SOX10^high^ melanoma cells. Conversely, the U1 ecosystem, associated with metastatic relapse, was characterized by undifferentiated NGFR^high^, ZEB1^high^, and SOX10^high^ – melanoma cells, colocalized with PDL1^-^ CD163 “M2-like” TAM and CD8^+^PD1^-^ T cells. These findings add to the growing body of evidence supporting coevolution of intra-tumoral heterogeneity and immune cell polarization (Plaschka et al. 2022; Barkley et al. 2022; Liu et al. 2024; H. Chen et al. 2025).

Importantly, the association of the U1 ecosystem in the primary tumour with macroscopic metastatic relapse suggests a role of this ecosystem in fostering metastatic founder cells (MFCs). This hypothesis is supported by recent data in mice demonstrating that in the primary tumours, melanoma cells with an undifferentiated/mesenchymal-like cell state can act as a reservoir of MFCs (Karras et al. 2022). Interestingly, recent data on human sentinel lymph node biopsies revealed that early disseminated cancer cells (DCC) entering the lymph node undergo a switch from a transitory phenotype to a more NCSC state upon exposure to interferon (Guetter et al. 2025). Thus, heterotypic tumour-immune ecosystems identified here might also foster the outgrowth of DCC towards overt metastases.

In summary, our study suggests that melanoma and immune cells co-evolve together in spatially restricted areas through cell-cell communications and integration of microenvironmental signals, forming diverse heterotypic ecosystems that can either enhance or limit the efficacy of adjuvant aPD1 ICI, ultimately influencing macroscopic relapse (Extended Data Fig.14). While spatial transcriptomic approaches offered valuable insights into the molecular pathways underlying these ecosystems, future mechanistic studies are required to elucidate their developmental origins and functional interdependencies.

## AKNOWLEDGEMENTS

The authors would like to thank Brigitte Manship for critical reading, the “Centre de Ressources Biologiques” from the “Hôpital Lyon Sud/Hospices Civils de Lyon” hospital for providing the tumour materials, Nicolas Gadot and Elodie Voilin from the Research pathology platform East, Cyril Deglétagne, Carole Audoynaud and Julie Valentin from the Cancer genomics platform (CGP). This work was funded by the Ligue Nationale contre le Cancer (Comité de l’Ain), the Lyon Integrated Research Institute in Cancer (SIRIC LYriCAN INCa-DGOS-Inserm_12563 and LYriCAN+ INCa-DGOS-INSERM-ITMO cancer_18003), the Institut Convergence PLAsCAN (ANR-17-CONV-0002), the ERiCAN program of Fondation MSD-Avenir (Reference DS-2018-0015), the Institut National contre le Cancer (INCA-DGOS PRTK), ARC sign’it 2019 Birdman, the IRICE Région Auvergne Rhône-Alpes grant for mIF, the Société Française de Dermatologie (SFD), the association Melarnaud and Vaincre le Mélanome. F.P. was supported by the Fondation pour la Recherche Médicale, grant number FDM202306017085, by “Année-Recherche” from the Agence Régionale de Santé Auvergne-Rhône-Alpes and by the “Prix Hervé Fridman”. M.D. was supported by a fellowship from Ligue Nationale contre le Cancer. F.B. was supported by a fellowship from ITMO Cancer of Aviesan within the framework of the 2021-2030 Cancer Control Strategy, on funds administered by Inserm. S.Du. was supported by a fellowship from the Association pour la Recherche contre le Cancer (ARC).

## AUTHOR CONTRIBUTION

J.C., S.Da., and A.E. conceived and jointly supervised the study. A.E., S.Da. and M.Do. contributed to sample and patient information collection. MDo performed pathological analyses. M.G., V.B., A.L., J.B., A.D. and F.P. performed mIF staining and image analysis. R.S. and L.T. performed QC for RNAseq, snRNAseq and spatial transcriptomic analyses and supervised bioinformatic analyses. F.P. and M.Du. performed bio-informatic analyses. A.L., B.D., A.D., C.C., J.V., A.E., J.L., S.Du., F.B., S.Da. and J.C. assisted with data analysis. F.P., M.Du., V.B. and A.L. analysed data and generated figures and tables for the manuscript. F.P., M.Du., A.L., J.V., J.B., B.D., C.C., S.Du., F.B., J.L., A.E., S.Da. and J.C. contributed to data interpretation. F.P., M.Du., A.E. and J.C. wrote and revised the manuscript, and all co-authors reviewed the manuscript.

## METHODS

### Clinical data of the patient cohorts

The cohort consisted of 57 pretreatment, (FFPE) sections of the entire primary melanoma from 57 patients with resected stage III (American Joint Commission on Cancer, 8^th^ edition [AJCC-8]) melanoma from the Lyon Sud Hospital (Hospices Civils de Lyon, Lyon, France). All patients were treated with a wide local excision of the tumor bed, complete lymph node dissection (if needed), followed with adjuvant aPD1 ICI with either nivolumab (480 mg i.v. every 4 weeks) or pembrolizumab (400 mg i.v. every 6 weeks) for 12 months (between 2019 and 2020). Patients should not have received any ICI or treatment with MAPK inhibitors prior to the surgery. Patients were followed-up with cerebral, chest and abdominal computed tomography scans every 3 months for 3 years then every 6 months for 2 years, or until relapse, after the first aPD1 infusion. All patients were followed-up at least 2 years after the first infusion of immunotherapy. Patients were classified according to their 24-month relapse status. Relapse was defined by either: radiological recurrence, clinical progression according to the investigator or death, whatever came first. Summary of the clinical data of the patients is shown in Table 1. Written informed consent was obtained from each patient. The studies were conducted in accordance with recognized ethical guidelines (Declaration of Helsinki). This project was approved by the Ethical Commission of the Hospices Civils de Lyon and approved by the review board (Comité Scientifique et Éthique des Hospices Civils de LYON CSE-HCL – IRB 00013204, study n° 22_5680).

### Multiplexed immunofluorescence stainings

3-µm tissue sections were cut from formalin-fixed paraffin-embedded human melanoma specimens. Sections were stained with five 7-color panels using the OPAL™ technology (Akoya Biosciences, Marlborough, Massachusetts, USA) on a Leica Bond RX (Leica Biosystem, Wetzlar, Germany) on the Research pathology platform East (ANAPATH RECHERCHE). Sections were digitized with a Vectra Polaris scanner (Perkin Elmer, USA). The panels included melanoma cell plasticity panel (ZEB1, ZEB2, MITF, SOX10, SOX9, NGFR, DAPI), T cells panel (CD3, CD8, SOX10, ZEB1, PD1, KI67, DAPI), TAM panel (CD68, pSTAT1, CD163, PD-L1, SOX10, DAPI), DC panel (CLEC9A, CLEC10A, LAMP3, BDCA2, CD8, SOX10, DAPI) and TLS panel (CD3, CD20, IgA, IgG, LAMP3, SOX10, DAPI). Details of the antibodies are in Extended Data Table 1. After staining, the slides were digitized with Vectra Polaris® (Perkin Elmer, Waltham, Massachusetts, USA).

### Multiplex immunofluorescence image processing and cell segmentation

Visualization and annotations of the regions of interest of each scan were performed with Phenochart® version 1.0.9 (Akoya Biosciences, Marlborough, Massachusetts, USA). Quality control of the images was performed by: slides with poor quality (n=4), heavily pigmented lesions precluding any immunostaining (n=2) were excluded. Segmentation of the scans was performed with inForm® version 2.4.8 (Akoya Biosciences, Marlborough, Massachusetts, US). The scans were then analysed with HALO-AI® image analysis software version 3.5 (Indica Labs Inc., London, UK).

Under HALO-AI®, tissue and nuclear segmentation were performed using the MiniNet module and the NucSeg FL 1.0.0 module, respectively. The slides were manually annotated: the first class was the “tumour”, and the second class the “stroma”. The “Analysis” module “FISH-IF 1.0.0” was used with the positivity threshold for the MFI of each OPAL™ channel set depending on its nuclear or cytoplasmic/membrane affinity and on the visual control. The resulting output was a .csv file per slide containing all the recognized cells.

### Fusion and informatic reconstruction of the three panels under HALO-AI®

The three immunofluorescence images were reconstructed under HALO-AI®. The images were then registered and fused with the “Serial qptiff fusion” module. Next, the fusion was performed by HALO-AI® that applied deformation and translation to match the three images. The .csv files were extracted as outputs.

### RNA sequencing analyses

RNA extractions were sequenced on the CRCL Cancer Genomics platform, on an Illumina NovaSeq machine with a paired-end protocol (64M reads). Libraries were prepared with the RNAexome FFPE kit from Illumina. Raw sequencing reads were aligned on the human genome (GRCh38) with STAR (v2.7.8a), with the annotation of known genes from gencode v37. Gene expression was quantified using Salmon (1.4.0) and the annotation of protein coding genes from gencode v37. Differential gene expression analysis, between immunotypes 1 and 3, was performed using DESeq2 (v 1.42.1). Genes that had a log2FoldChange> 1 and an adjusted pvalue (padj) <0.05 were kept for further analysis. The volcanoplot was realised using ggplot2. Then, Gene Set Enrichment Analysis (GSEA) was performed using the enrichGo function from the clusterProfiler package (v 4.10.1), to identify enriched GO terms based on the identified DEGs. Finally, pathways enrichment results were sorted based on geneRatio and visualized with barplots using ggplot2.

### Visium data generation and preprocessing

Spatial transcriptomic data were generated by the CRCL Cancer Genomics Platform using Visium CytAssist Spatial gene expression for FFPE, Human Transcriptome, kit (10X Genomics) according to the manufacturer’s guidelines. Briefly, five micron FFPE section were placed on histological slides (TOMO), deparaffinized, stained with hematoxylin and eosin (H&E) by the Research Pathology Platform at CRCL. After imaging, 6.5 x 6.5 mm areas were selected in the tissue section for each patient. After destaining and decrosslinking steps, human Visium probes were hybridized on the samples, ligated and transferred on the Visium capture slides using the CytAssist instrument (10X Genomics). Libraries were then prepared and sequenced on a NovaSeq 6000 sequencer (Illumina) targeting 50,000 reads/spot. Visium data were then processed using SpaceRanger from 10X Genomics (v 3.0.0), with the "count" algorithm and the human genome reference refdata-gex-GRCh38-2020-A for quantification. Transcript capture was performed using the Visium_Human_Transcriptome_Probe_Set_v2.0_GRCh38-2020-A.csv probeset from 10X Genomics.

### Identification of tumour-enriched spots

Spot labelling was performed using 1) histopathological reviews by our pathologist specialized in dermatopathology and 2) with the minimal lineage gene (MLG) signature described by Pozniak et al, which is designed to discriminate melanoma cells from cancer-associated fibroblasts. The AddModuleScore (Seurat v5.1.0) function was applied with a threshold of 4 to classify spots: an MLG score > 4 indicated tumour enrichment of the spot, while an MLG score <4 indicated stromal enrichment of the spot. To further characterize the tumoral spots, the normalized expression of MITF was used to label them as MITF^high^ (MITF normalized expression > 1.8) and MITF^low^ (MITF normalized expression < 1.5). To access the differentially expressed genes across spots, the FindAllMarkers function was applied, with the following thresholds: avg_log2FC > 1 and pval_adj < 0.05. The tool “What is my melanoma signature” (https://wimms.tanlab.org/)(Hu et al. 2024) was used to generate the radar plots, based on the identified DEGs, allowing to characterize the spots based on public gene signatures. GSEA was also performed on those same DEGs, using the fgsea function from the fgsea package (v 1.28.0) and the hallmark and GOBP database from the package msigdbr (v 7.5.1). GOBP pathways were visualized on the Visium slides using AddModuleScore, the SpatialFeaturePlot function from the Seurat package and ggplot2 (v 3.5.1)

### snRNAseq data generation and preprocessing

The generation of snRNAseq dataset was performed following the snPATHOseq protocol from Wang et al 2024 (Wang et al. 2024), using FFPE tissue sample from 2 treatment-naïve cutaneous metastases. Libraries were prepared using the Next GEM Flex protocol. SnRNAseq data were processed through CellRanger from 10X Genomics (v 7.1.0), using the "multi" algorithm and the human reference genome refdata-gex-GRCh38-2020-A for quantification. Transcript capture was performed using the Chromium_Human_Transcriptome_Probe_Set_v1.0.1_GRCh38-2020-A.csv probeset from 10X Genomics.

### Cellular deconvolution of the spots

To constitute an in-house reference dataset, snRNAseq data from the two treatment-naïve cutaneous metastases were merged. Quality control filtering was performed using the following thresholds: nFeature_RNA >150, nCount_RNA < 150000, percent.mt <20. Data were normalized using SCTransform (v 0.4.1), regressed on cell cycle and nFeature_RNA. Then, Principal Component Analysis (PCA) was performed using runPCA. Batch effect was corrected using harmony (v 1.2.1). Clusters were defined using FindNeighbors and Findclusters. Finally, A Uniform Manifold Approximation and Projection (UMAP) was computed through RunUMAP. Doublets were removed using Scrublet (v 0.2.3) on Python. The whole TME was largely annotated, using signature and gene markers from the literature. The myeloid compartment was then isolated and reclustered using the same pipeline, and TAM signatures (R.-Y. Ma, Black, et Qian 2022) were used to finely annotate this compartment. CARD (v 1.1) package (Y. Ma et Zhou 2022) was used to perform cellular deconvolution of the spots, using the snRNAseq dataset composed of 2 cutaneous metastases as reference.

### Spots neighbourhood analyses

Spots neighbourhoods were defined based on the specific hexagonal geometry of Visium spots. A spot was considered to be within the neighbourhood of a tumour spot if it was located within the first 2 rows (corresponding to an approximate radius of 200 µm) around the spots of interest and labelled as stroma enriched. Spots that were simultaneously present in the list of differentiated and undifferentiated spots neighbourhood were labelled as mixed-neighbourhing non tumoral spots (Fig.3g). The spots were CLR normalized based on the deconvoluted cellular fraction of TAM clusters, clustered using Seurat, and annotated according to the predominant TAM subtypes. The “IFN TAM & C1QH TAM”, “Angio-TAM” and “MoMacs” spots were further kept for abundance analyses between differentiated versus undifferentiated spots using the miloR package (v 2.0.0), using a FDR threshold of 0.1 to account for multiple comparisons (Dann et al. 2022).

### Cellchat analysis between tumour and TAM-enriched spots

TAM-enriched spots were identified using the AddmoduleScore function and a macrophage gene signature derived from MCPcounter signatures. Stroma enriched spots with a macrophage score > 0.5 were classified as TAM-enriched. Next, adjacent TAM-enriched spots were defined as spots located within the neighbourhood of differentiated or undifferentiated tumour spots (as described above), contrary to distant TAM-enriched spots. Cell-cell interactions were inferred using CellChat package (v 2.1.2), with Contact = FALSE and a fixed distance of 200 µm, between differentiated/undifferentiated spots and TAM enriched spots within this spatial range. Interactions that were conserved were the one that were either unique to one condition (Differentiated or Undifferentiated), or with a significantly higher probability in one condition compared to the other. Chord plots representing these interactions were assessed using the CellChat and the circlize (v 0.4.16) packages.

### pSTAT1, Interferon Gamma Response and hypoxia scoring

The pSTAT1 gene signature was scored among macrophages subset using AddModuleScore. The ViolinPlot was visualized using VlnPlot2 from the SeuratExtend package. This pSTAT1 signature was defined as a combination of 3 signatures (Rodero et al. 2017; Scott et al. 2021; Adang et al. 2024) (Extended Data Table 2). Interferon Gamma Response signature was extracted from hallmark database. The hypoxia signature was the one described in Di Giovannantonio et al. (Di Giovannantonio et al. 2025).

All patients’ slides were annotated to identify differentiated and undifferentiated spots as described above. Then, all patients’ slides were merged and SCTransform was applied to normalize the data. AddModule score was used to compute a score for each signature or pathway. Then a Mann-Whitney U test was performed on each score, between differentiated and undifferentiated spots, and boxviolinplots were used for visualization purposes, with the ggplot2 package.

### Cut-off’s settlement

Every quantitative variables analysed with Log-rank test were converted into a binary variable (high/low) according to a cut-off value automatically defined with Cutoff Finder v1(Budczies et al. 2012). The method for cut-off determination was the Euclidean distance.

### Multi-IF spatial analyses

#### K-means clustering of tumour cells

Every SOX10+ tumour cells across all samples were retrieved into a single .csv file containing the MFI of each phenotypic marker. Then, a k-means clustering was performed. The optimal number of clusters was determined using the silhouette score.

### Melanoma cell states patches detection

For each tumour, the .csv file containing the spatial coordinates and the phenotypes of every tumour cells was converted into a SpatialExperiment object. Next, a spatial interaction graph was built (‘buildSpatialGraph’ function) with a radius of 25 µm with the imcRtools R package (v 1.10.0)(Windhager et al. 2023). The ‘patchDetection’ function was then run with a minimal number of cells of 5. This operation returned the SpatialExperiment object with a new column containing the patch identifier for each cell for a given melanoma cell state.

### Spatial analyses of immune cells

For the clustering of DCs, the ‘FindNeighbor’ function from the SPIAT package (v 1.6.2) was used setting the minimal number of cells to form a cluster to 2 and the radius to 50 µm. An aggregate was defined as at least 10 DCs within a radius of 50 µm. The proximity score corresponded to the normalized mixing score (NMS) from the SPIAT package and was the main spatial metric used to account for the baseline density of both tested immune cells(Feng et al. 2023). A radius of 25 µm was chosen for NMS analyses.

### Immunotypes identification

First, the patients for whom TAM and T cells parameters were available were retained (48 out of 57 patients). Next, every TAM and T cell densities significantly associated with RFS were included in the Partitioning Around Medoids clustering of patients (a non-parametric k-means clustering). All features were transformed with the following operation: x -> log10(1e-3+x) and z-normalized across the dataset prior to inclusion in the model. A silhouette score was computed to select the optimal number of clusters.

#### Comparisons of the immune cell densities between melanoma cell states areas

From the fused images of the 8 selected tumours, homotypic differentiated (C2-MITF^high^, C3-SOX10^high^) and undifferentiated (C6-ZEB1^high^, C4-NGFR^high^) patches were manually contoured from the same tumours. Then, the area surface was calculated, the counts of TAM and T cells were extracted and the density of each immune subsets was computed within the different patches.

### Immune cellular neighbourhoods

From the fused images of the 45 tumours, melanoma patches of clusters C2-MITF^high^, C3-SOX10^high^, C4-NGFR^high^ and C6-ZEB1^high^ were retrieved and their border was extended by 50 µm with the function ‘findMilieu’ from the imcRtools R package (Windhager et al. 2023) to constitute immune CN. Briefly, the number of each TAM and T cell subsets was retrieved in each CN. Only CNs with a total number of TAM and T cells of at least 10 were kept. The area of each CN was calculated by considering them as concave polygons with the functions ‘concaveman’ from the concaveman package (v 1.1.0) and ‘st_polygon’ from the sf package (v 1.0.19). Subsequently, the count of each immune cells was divided by the area of the given CN, then by its corresponding frequency across the entire section of the given tumour to account for interpatient heterogeneity. Next, a Seurat object was built with every CNs from the 45 tumours and these CNs were clustered using Louvain function (resolution = 0.2) from the Seurat package (v 5.2.0). To study the enrichment of the generated CN clusters within specific melanoma cell states and to link it with ICI efficacy, the miloR package (v 2.0.0)(Dann et al. 2022) was used (k = 30, d = 9) with a subsampling rate of 10% and a FDR threshold set at 0.1.

### Entropy-based spatial analyses

The entropy-based analysis as proposed by Feng et al.(Feng et al. 2023) with the SPIAT automatically applied a fishnet approach to divide the tumour into n*n tiles (‘grid_metrics’ function, n = 20). For each individual tile, it returned a localized entropy that could then be compared to a threshold value. Next, a colocalization score was computed, consisting of the percentage of tiles with an entropy value above the threshold across the whole tumour. To dynamically determine this threshold for each cell types combination in order to account for the probability of these cell types to be colocalized by chance alone, and the intra- and inter-tumoral variation of the cell types density, we calculated the global entropy of a given tumour using the ‘calculate_entropy’ function and the returned value would then become the threshold value. We excluded empty tiles corresponding to the void around the tumours.

### Identification of the D1 ecosystem

The prevalence score of pairwise entropy-based interactions associated with an improved RFS (hazard ratio <1) with a P value < 0.10 were included in a co-occurrence matrix across each individual tumour to determine recurrent patterns of cell type interactions using the Jaccard’s similarity index from the ‘CooccurrenceAffinity’ package (v 1.0.0) (Mainali et Slud 2022). These interactions were then clustered based on this index with a Leiden clustering to identify highly recurrent pairwise colocalization patterns that might represent broader and more complex ecosystems. This led to the identification of the ecosystem D1.

### Regularized Cox regression analyses

Cox proportional hazard models were L1 (LASSO or Least Absolute Shrinkage and Selection Operator) regularized logistic regression models, fitted using the glmnet R package (v 4.1.8) (Friedman et al., 2010). Briefly, such approach enables the selection of variables with the strongest relationship to the outcome and decreases the risk of overfitting. For the MPR, features were computed under the transformation x - > log(1e-3+ x) and z-normalized across the dataset prior to inclusion in any models.

For the calculation of the composite score, the MPR, the immunotypes and the ecosystems were included in the Cox regression analyses. Regarding variable pre-processing, the MPR was binarized according to its cut-off value (“low” vs “high”). The immunotypes corresponded to the cluster number as described before (“1”, “2” or “3”, Fig.2b). The ecosystem variable was the scenario number (Fig.6e) depending on the U1 or D1 ecosystems prevalence (“1”, “2”, “3”, “4”).

Next, multicollinearity was measured with a variance inflation factor (vif) on a Cox proportional hazard model including the MPR, the immunotypes, the ecosystems and their interaction terms. Every vif value were equal to 1, meaning that no collinearity was detected. Afterwards, the Cox model was fitted using L1 penalty. A cross validated error plot based on the Harrell’s C index could then help in selecting the optimal λ value where the curve hits its minimum. Such λ value was used in the finalized fitted Cox model. From this model, we could extract the coefficients (w) associated with each of the *n* included variables (v) and calculate for each patient the composite score as followed:

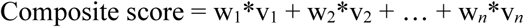

Of note, in such models, positive coefficients are associated with worse RFS whereas negative coefficients are associated with better RFS.

### Comparisons between Cox proportional hazard models

For each clinical, histological or biological prognostic variable, a univariate Cox proportional hazard model was computed and the Harrell’s C index was compared with that of a univariate model based on the Breslow index, which is the best prognostic variable in clinical practice. The ‘compareC’ function from the compareC R package (v 1.3.2) was used.

### ROC curve analyses

When performed, Receiver Operating Characteristic (ROC) curves were generated with the pROC R package (v 1.18.5) using the function ‘roc’, with respect to predict the 2-year recurrence status. The area under the curve (AUC) was calculated with the function ‘auc’.

### Survival analyses

Univariate Cox proportional hazard model were fitted for the tested parameters and a Log-rank test was performed to compare the different groups, in respect to RFS. The time to event was computed as the difference in months between the date at diagnosis of the primary melanoma and the date of the first recurrence or death, whatever came first. For RF patients, this date corresponded to the date of last survival update.

### Data availability

The data reported in this paper have been deposited in the Gene Expression Omnibus (GEO) database under accession number GSE300445 and GSE300446.

Public single-cell RNAseq (Pozniak et al. 2024) data of human metastatic melanoma were retrieved from the KU Leuven Research Data Repository.

### Statistical analysis and plotting

An α risk of 5% was applied for every statistical analysis. Statistical analyses were performed using RStudio with R. Statistical tests used included: non-parametric Mann-Whitney U tests for unpaired 2-group comparisons, paired Wilcoxon tests for paired 2-group comparisons, Log-rank tests for survival analyses, non-parametric Kruskal-Wallis tests for 3 or more unpaired groups, and Friedman test for 3 or more paired groups with post-test between-group comparisons performed with paired Wilcoxon tests, which P values were adjusted for multiple comparisons with the ‘Benjamini-Hojberg’ method. For correlation tests, a non-parametric Spearman test was utilized. Regarding boxplots, the whiskers extend to the minimum and maximum values within 1.5 times the inter-quartile range from the 25^th^ and 75^th^ percentiles.

---

**Extended Data Figure 1 (related to Fig.1):**
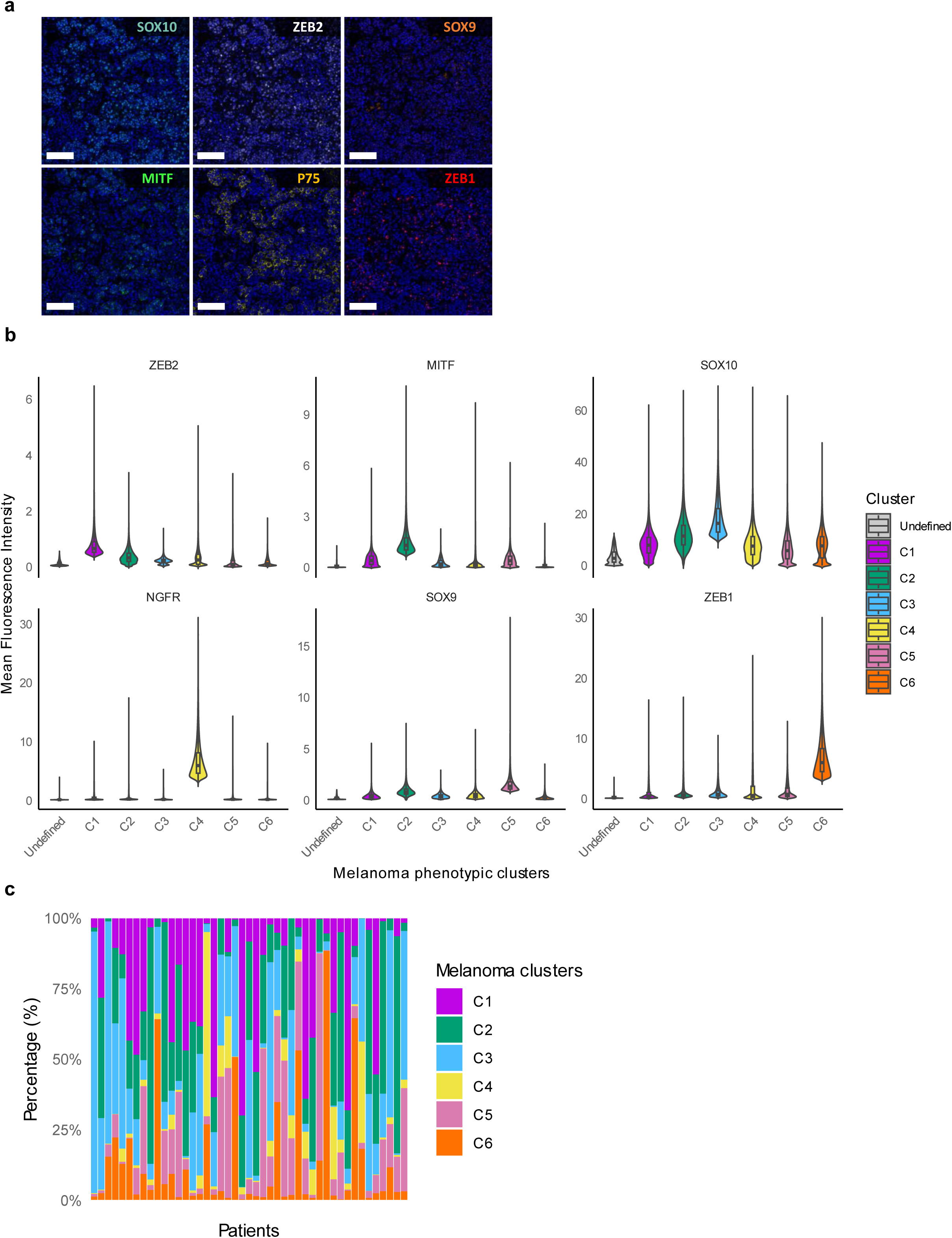
Identification and quantification of melanoma cell states within primary tumours. **a** Detailed mIF panel for the study of differentiated and undifferentiated melanoma cells. DAPI counterstaining appears as blue within each panel. Scale bar = 100 µM. **b** Mean fluorescence intensity (MFI) of the six markers used in tumour cells according to phenotypic clusters defined in (**Fig.1c**). **c** Stacked bar plots of C1, C2, C3, C4, C5, C6 clusters within each tumour, showing all clusters per patient (*n* = 45).

**Extended Data Figure 2 (related to Fig.1):**
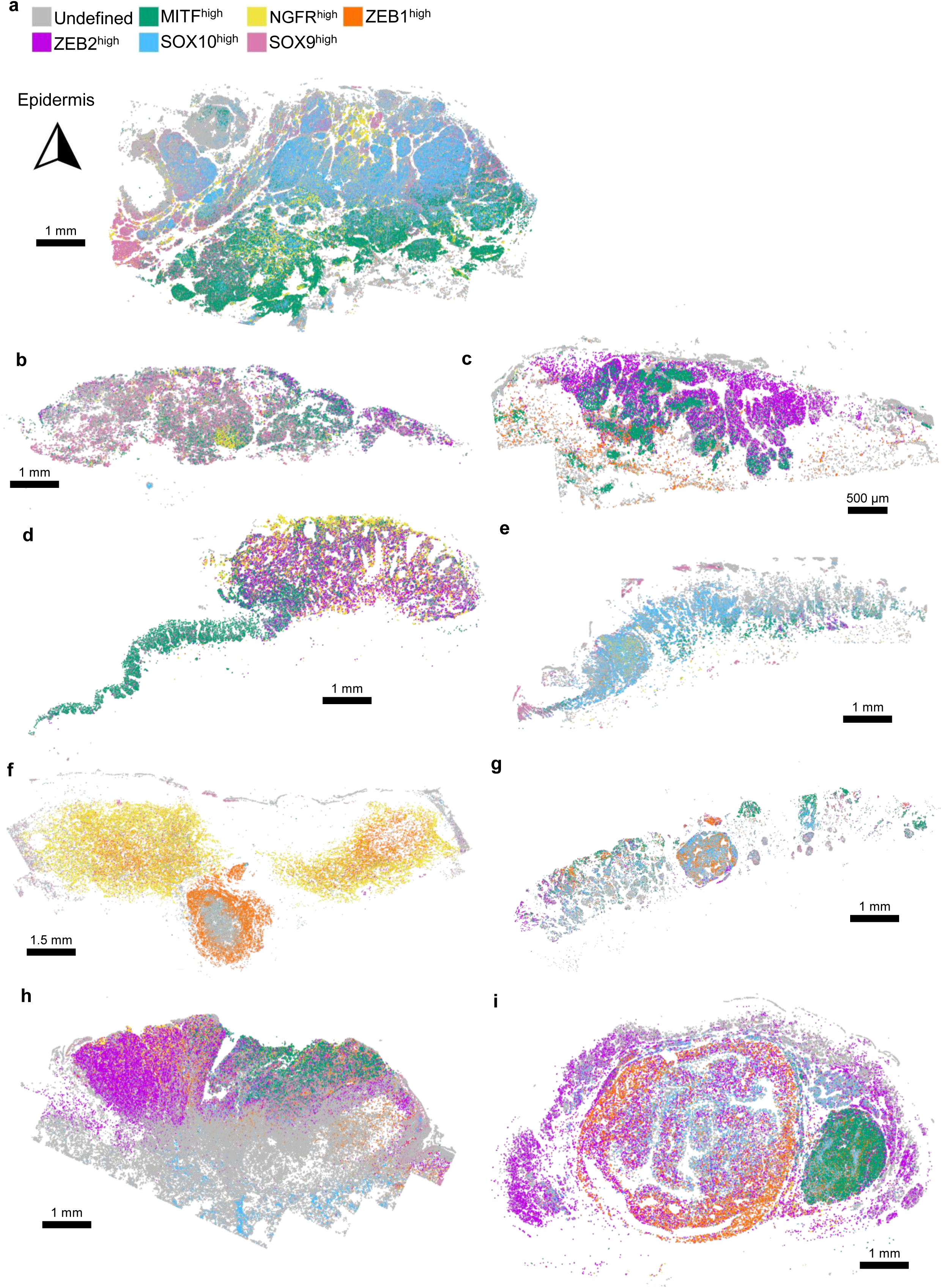
Informatic reconstruction of intra-tumoral and inter-tumoral heterogeneity. **a-i** Nine representative primary melanoma tumour reconstructions (of n = 45), illustrating the phenotypic inter-tumoral and intra-tumoral heterogeneity. In each example, the epidermis is located at the upper part of the tumour.

**Extended Data Figure 3 (related to Fig.1):**
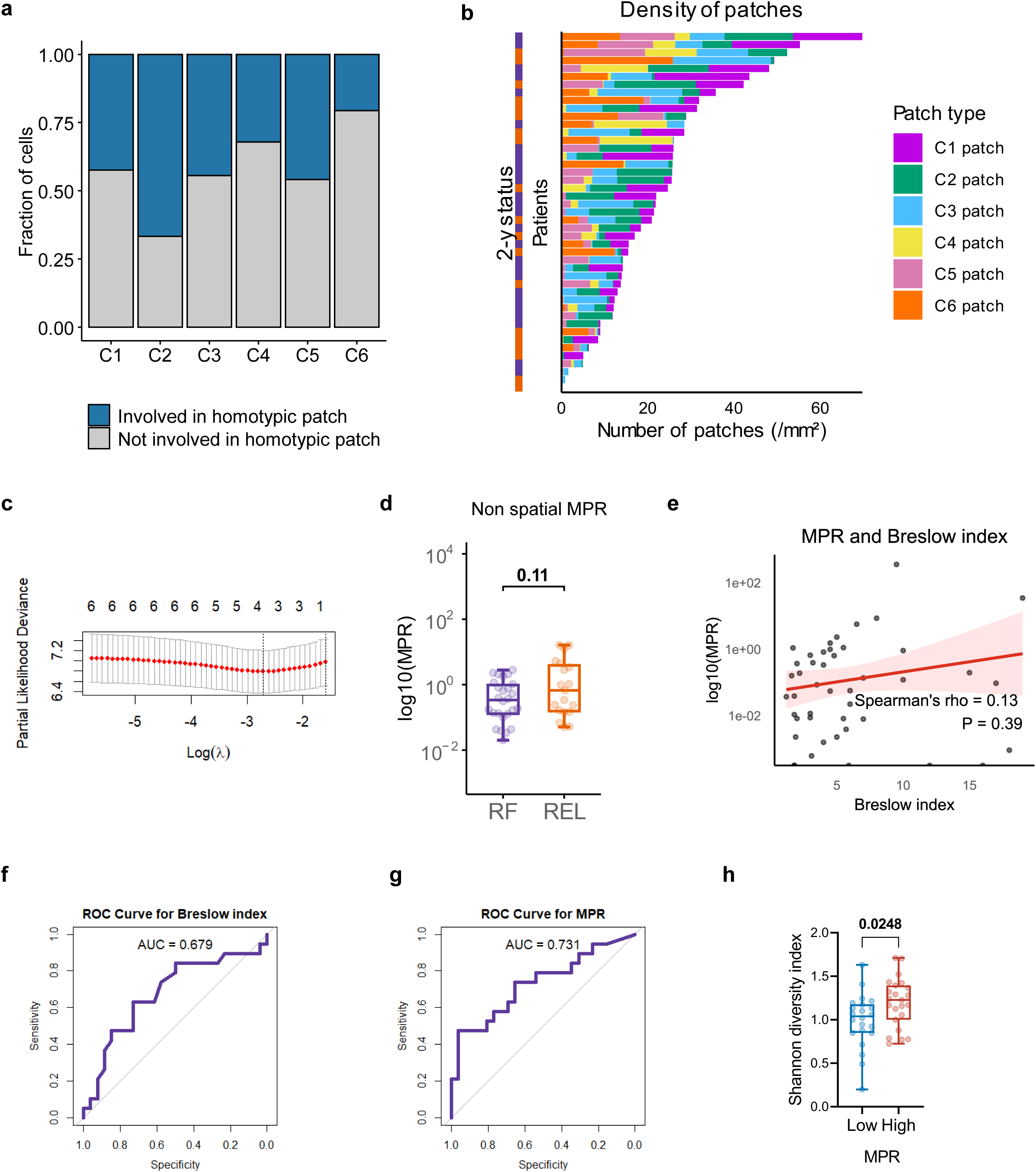
Spatial heterogeneity of melanoma tumours and the melanoma plasticity ratio. **a** Fraction of melanoma cells involved (blue) or not (in gray) in homotypic patches according to phenotypic clusters. **b** Stacked bar plots of melanoma phenotypic clusters representing the proportion of melanoma cells involved in a homotypic patch. Each row represents a patient. **c** Lasso partial likelihood deviance of the phenotypic clusters. The dotted line indicates the minimal lambda value that was retained for the final Cox model. **d** The melanoma plasticity ratio (MPR) based on all C4+C6 against C2+C3 tumour cells, irrespective of their participation in homotypic patches. Mann-Whitney U test between RF (purple) and REL (orange) patients, the p-value is indicated. **e** Correlation plot between the MPR and the Breslow index. The 95% confidence interval is filled in light red.’ test. The p-value is indicated. **f** Receiver Operator Curve (ROC) for the Breslow Index to predict the 2-y status. The Area Under Curve (AUC) value is plotted beside the ROC curve. **g** ROC for the MPR to predict the 2-y status. The AUC value is plotted beside the ROC curve. **h** Shannon diversity index between between MPR^high^ (red) and MPR^low^ (blue) patients.

**Extended Data Figure 4 (related to Fig.2):**
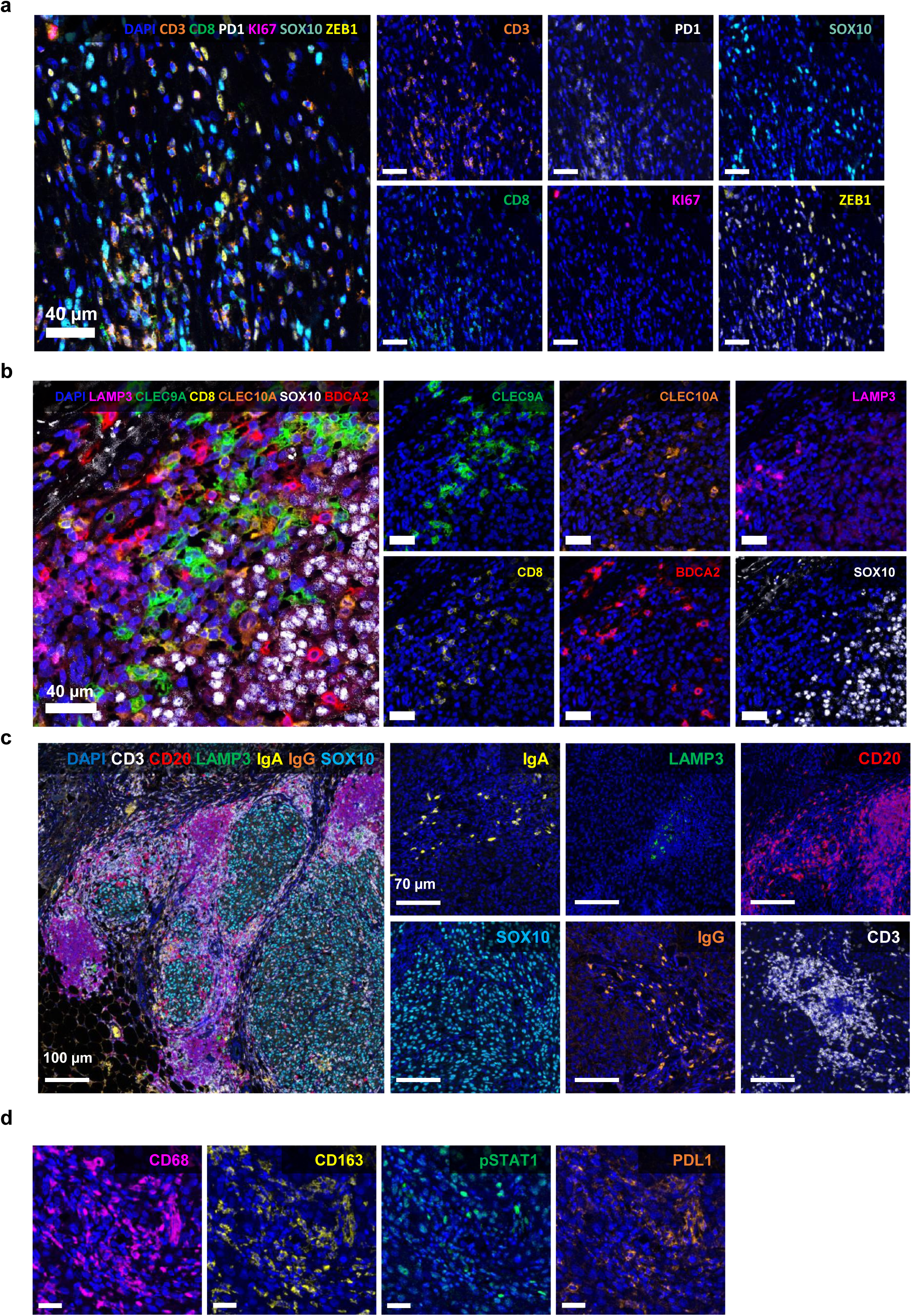
Multiplex immunofluorescence (mIF) panels used to study immune cells. **a** Representative tissue (of *n* = 57 tumours) section showing the mIF markers used to study T cells. Scale bar = 40 µm. **b** Representative tissue (of *n* = 43 tumours) section showing the mIF markers used to study dendritic cells. Scale bar = 40 µm. **c** Representative tissue (of *n* = 57 tumours) section showing the mIF markers used to study tertiary lymphoid structures. Left scale bar = 100 µm and scale bars from the small images = 70 µm. **d** Representative tissue (of *n* = 48 tumours) section showing the mIF markers used to study tumour-associated macrophages. Scale bar = 40 µm.

**Extended Data Figure 5 (related to Fig.2):**
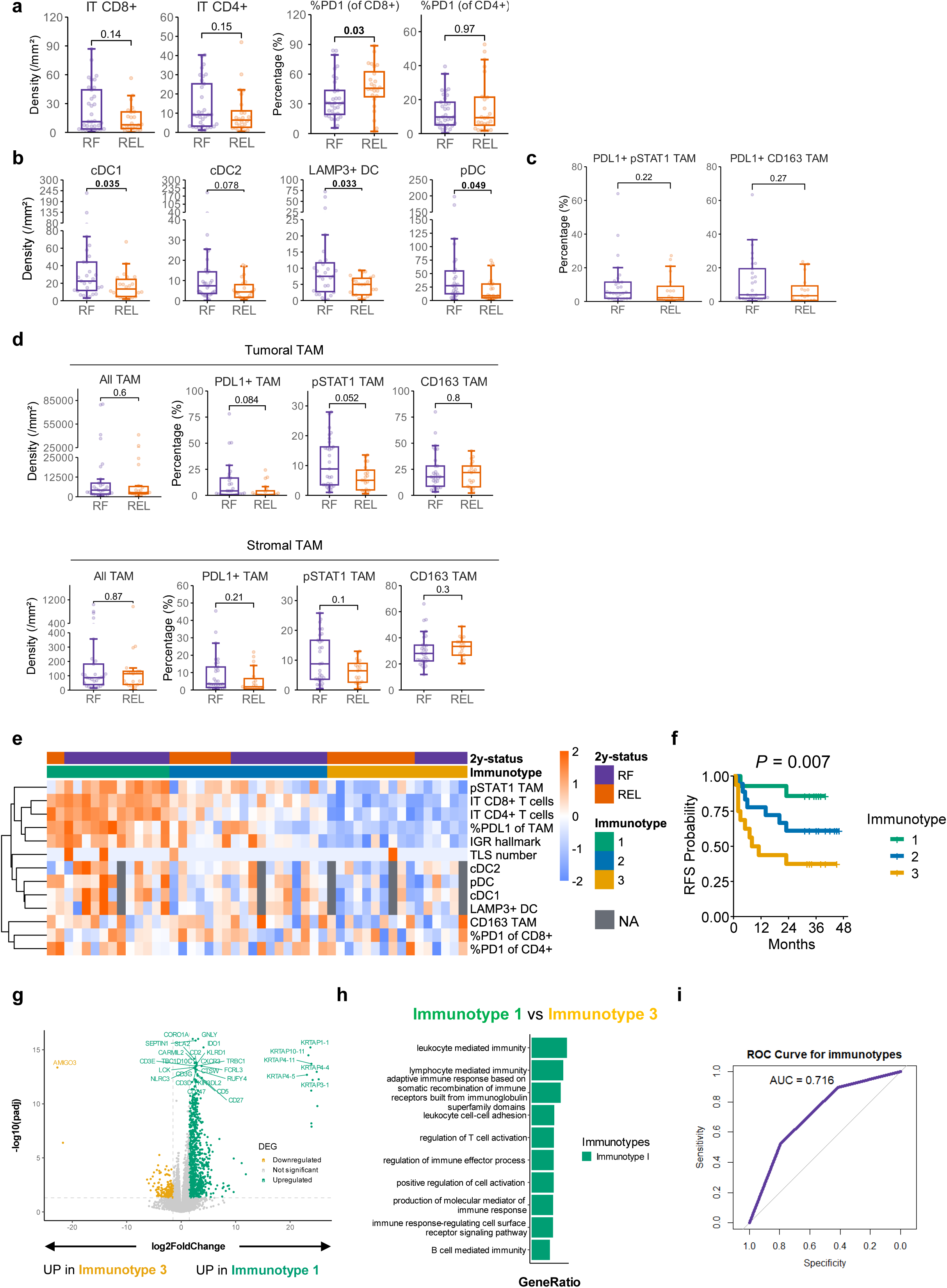
Integrative analyses of T cells, dendritic cells and tumour-associated macrophages densities. **a** Density and proportion of T cell subsets, according to 2-y status (*n* = 57). Mann-Whitney U tests, p-values are indicated on the boxplots. IT: intra-tumoral. **b** Density and proportion of dendritic cells subsets, according to 2-y status (*n* = 42). Mann-Whitney U tests, p-values are indicated on the boxplots. **c** PDL1^+^pSTAT1 TAM expressed as percentage of all pSTAT1 TAM (left) and PDL1^+^CD163 TAM expressed as percentage of all CD163 TAM (right). Boxplots according to the 2-y status (*n* = 48). Mann-Whitney U tests, p-values are indicated on the boxplots. **d** TAM density and proportions in the tumoral compartment (upper row) and in the stromal compartment (lower row), according to the 2-y status (*n* = 48). Mann-Whitney U tests, p-values are indicated on the boxplots. **e** Heatmap of immune cell frequencies from the mIF analyses, in addition to the IGR hallmark established by ssGSEA on RNA-seq data. Each column represents a patient (*n* = 48). Patients are clustered with Partition Around Medoids clustering into three immunotypes. DC: dendritic cell, IGR: interferon-gamma response, IT: intra-tumoral, NA: not available, REL: relapse, RF: relapse-free, ssGSEA: gene-centric single sample Gene Set Enrichment Analysis, TAM: tumour-associated macrophage, TLS: tertiary lymphoid structure, y: year. **f** Kaplan-Meier curves of relapse-free survival (RFS) according to the immunotypes obtained in (e). Log-rank test, the p-value is indicated on the graph. **g** Volcano-plot of the variable genes in RNA-seq between patients with immunotype 1 versus 3. DEG: differentially expressed gene. **h** Gene Set Enrichment Analysis (GSEA) of enriched pathways from DEGs between patients with immunotype 1 versus 3. **i** Receiving Operator Curve (in purple) of the immunotypes according to the 2-y status. The Area Under Curve (AUC) value is plotted beside the curve.

**Extended Data Figure 6 (related to Fig.2):**
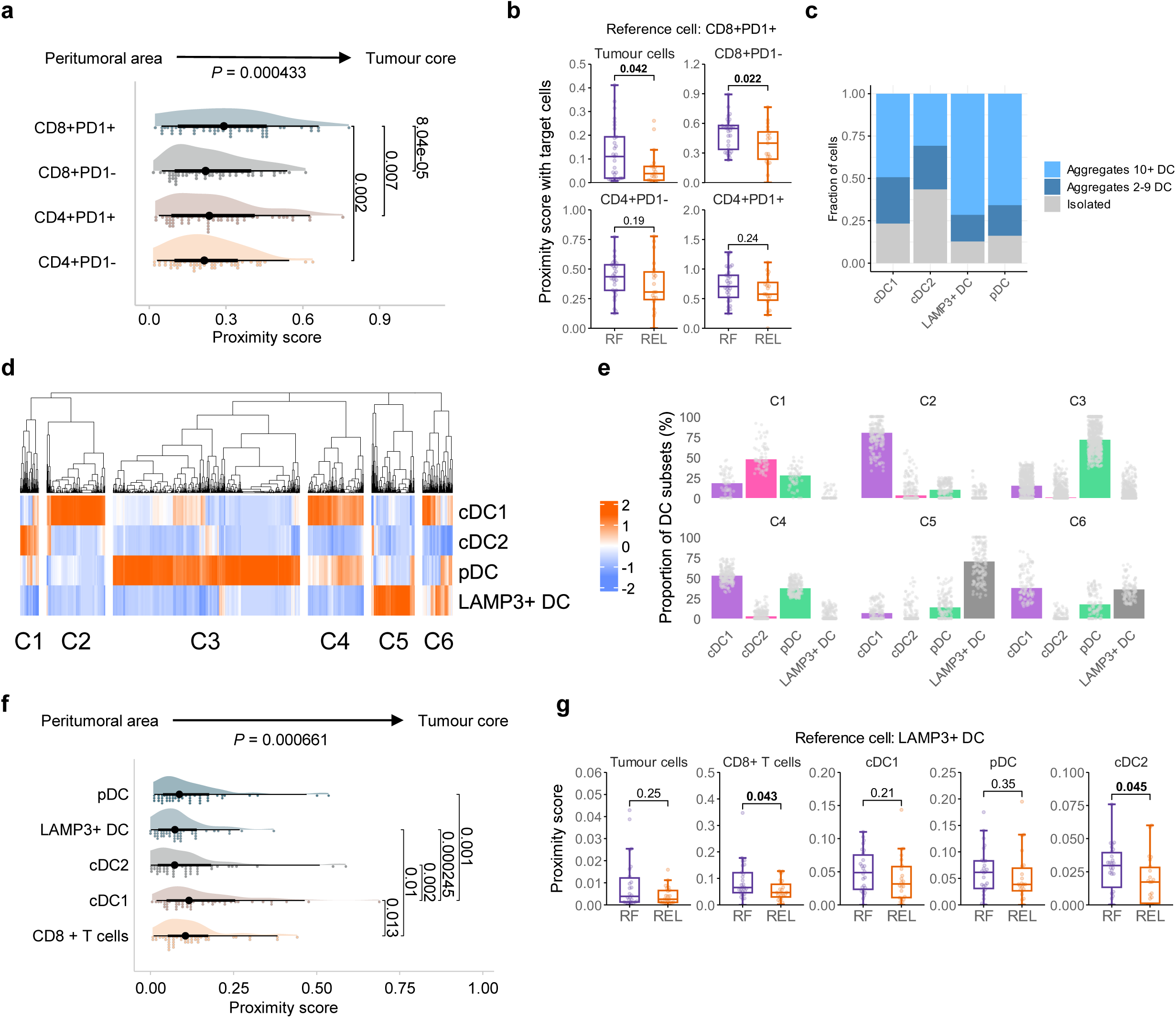
Spatial profiling of T cells and dendritic cells. **a** Proximity score illustrating the intra-tumoral infiltration by T cells subsets in a perimeter of 25 µm around a tumor cell (*n* = 57). A higher score indicates more colocalization between both cell types independently of their frequencies. Friedman test and paired Wilcoxon tests, with p-values adjusted by Benjamini-Hochberg method. b Proximity score between CD8+PD1+ T cells and target cells within a radius of 25 µm, according to the 2-y status (*n* = 57). Mann-Whitney U tests, p-values are indicated on the boxplots. **c** Stacked bar plot showing the types of aggregates according to the DC subsets (*n* = 42). **d** Heatmap showing DCs spatial aggregation patterns into 6 clusters (from C1 to C6). **e** Bar plots representing the composition of the DCs clusters identified in (**d**). Each dot represent a DC. **f** Proximity score illustrating the intra-tumoral infiltration by DC subsets in a perimeter of 25 µm around a tumour cell (*n* = 42). A higher score indicates more colocalization between both cell types independently of their frequencies. Friedman test and paired Wilcoxon tests, with p-values adjusted by Benjamini-Hochberg method. **g** Proximity score between LAMP3+ DC and tumour cells, CD8+ T cells, and other DC subsets according to the 2-y status (*n* = 42). Mann-Whitney U tests, p-values are indicated on the boxplots.

**Extended Data Figure 7 (related to Fig.3):**
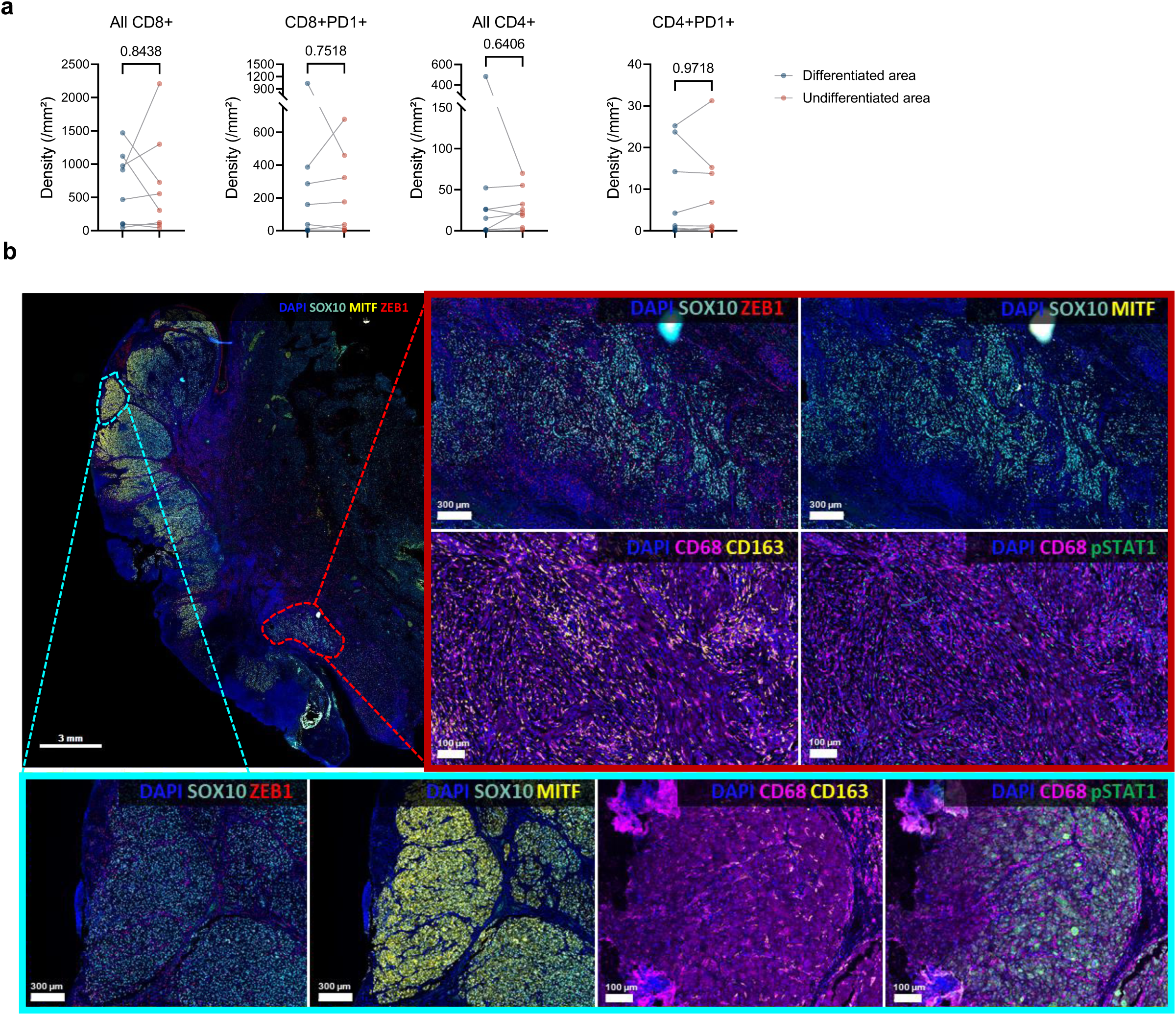
T cells and TAM subset analyses in areas containing differentiated and undifferentiated melanoma cells. **a** Paired comparisons of T cell subsets between differentiated and undifferentiated areas. Paired Wilcoxon tests, p-values are indicated on the boxplots. **b** Representative mIF image of (n = 45) TAM subsets according to the differentiated (SOX10+MITF+ in dashed blue) and the undifferentiated (SOX10+ZEB1+ in dashed red) areas within the same tumour. Scale bar = 3 mm. The red and blue frames are enlarged views of the undifferentiated, and differentiated areas, respectively.

**Extended Data Figure 8 (related to Fig. 3):**
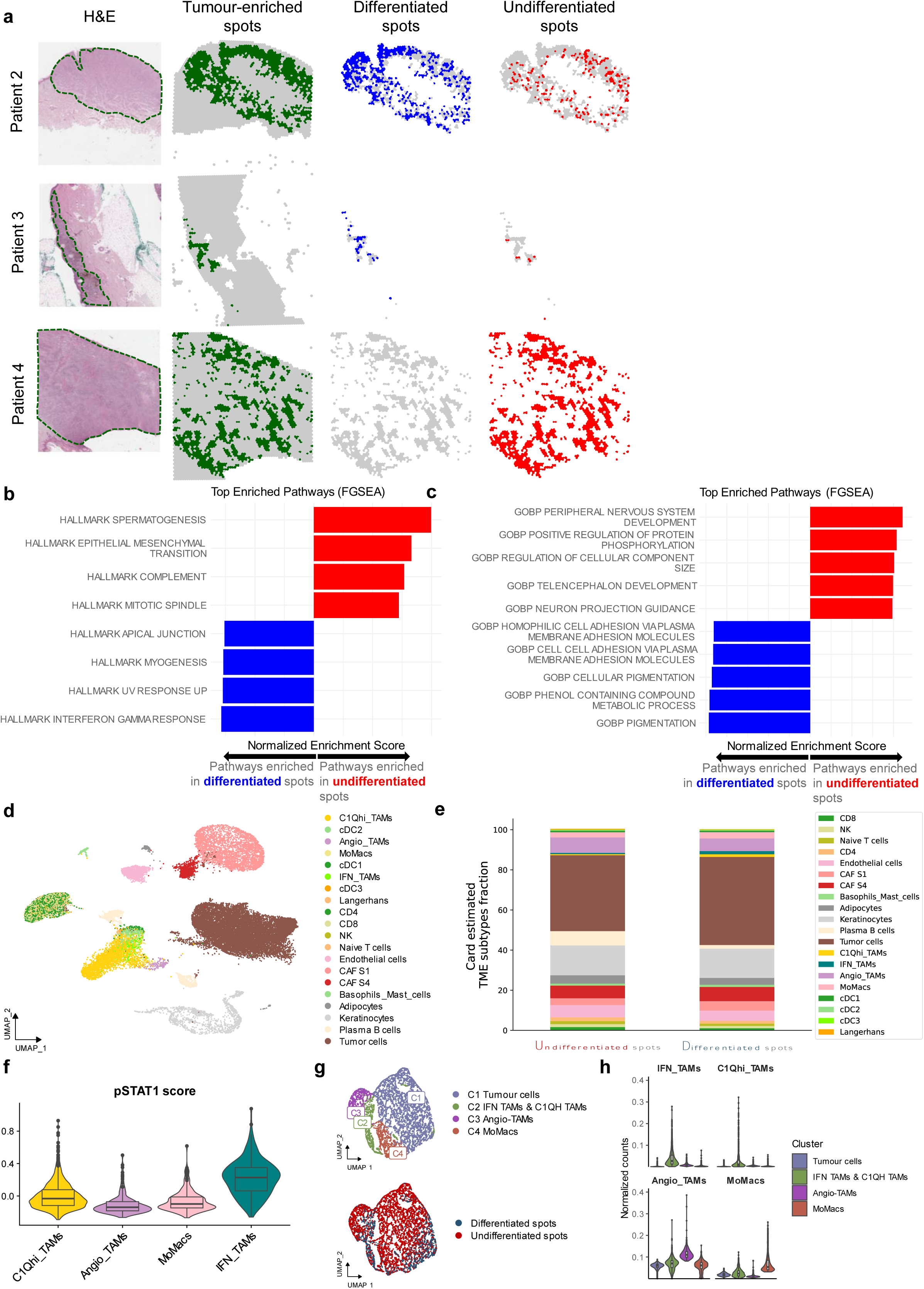
Spatial transcriptomics (Visium) analyses of TAM composition. **a** Identification of tumour-enriched spots: by pathologist annotation (first column, in dashed green), and by the binarized MLG signature (second column, in green). Identification of MITF^high^ spots (third column, in blue) and MITF^low^ spots (last column, in red), for patients #2, 3, 4. **b** Top enriched HALLMARK pathways by GSEA analysis on differentiated and undifferentiated spots, from all 4 patients (from **a** and Fig.3d). **c** Top enriched GOBP pathways by GSEA analysis on differentiated and undifferentiated spots, from all 4 patients. **d** Uniform Manifold Approximation and Projection (UMAP) of all the cell populations from the 2 snRNAseq metastases samples used for deconvolution. **e** Stacked bar plots showing the differences in proportion of deconvoluted cell types by CARD algorithm between undifferentiated and differentiated spots, among all 4 patients. **f** pSTAT1 score across TAM subsets from the 2 snRNAseq metastases dataset. **g** UMAP of the selected spots in (Fig.3g): annotations according to cellular composition (above), and according to the spot annotation in (Fig.3g) (below). **h** Violin boxplots of the normalized counts of TAM subsets in each cluster.

**Extended Data Figure 9 (related to Fig. 4):**
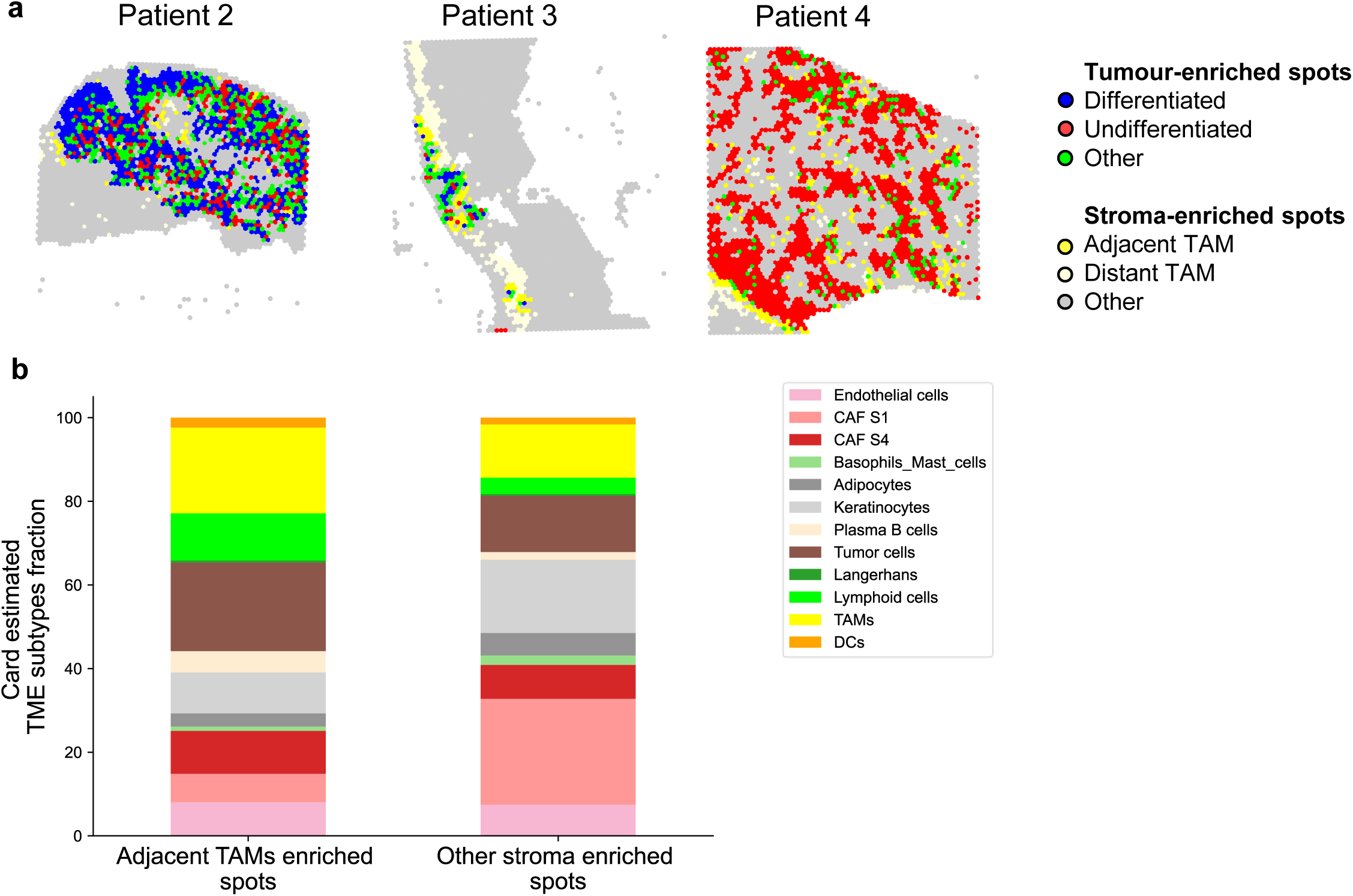
Pipeline for cell-cell communication analyses. **a** Labeling of cell spots for patients #2, 3 and 4 for cell-cell communication analyses, as depicted in Fig.4a. **b** Stacked bar plots showing the cellular deconvolution results in Adjacent TAM enriched spots (left) and Stroma enriched spots (right).

**Extended Data Figure 10 (related to Fig. 4):**
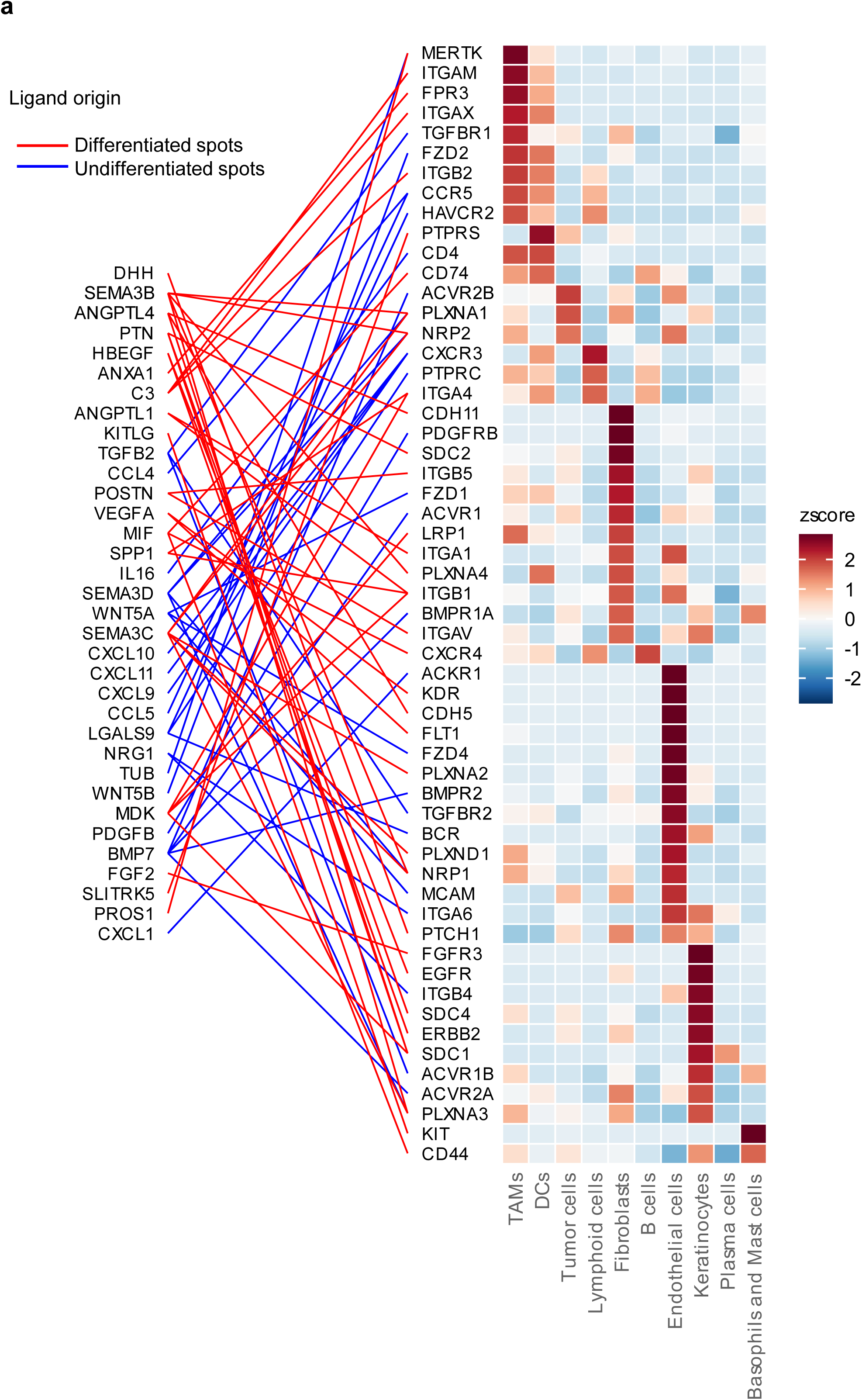
Ligands and receptors gene expression in a public single-cell RNAseq dataset. **a** Heatmap showing the gene expression as a z-score of the receptors from predicted cell-cell communications in (Fig.4b), among cellular clusters from the tumour microenvironment (right) in the Pozniak et al (2024) dataset. The lines represent the ligand-receptor(s) interactions and are coloured according to the tumour-enriched spot of origin of the ligands (differentiated in blue, and undifferentiated in red).

**Extended Data Figure 11 (related to Fig. 4):**
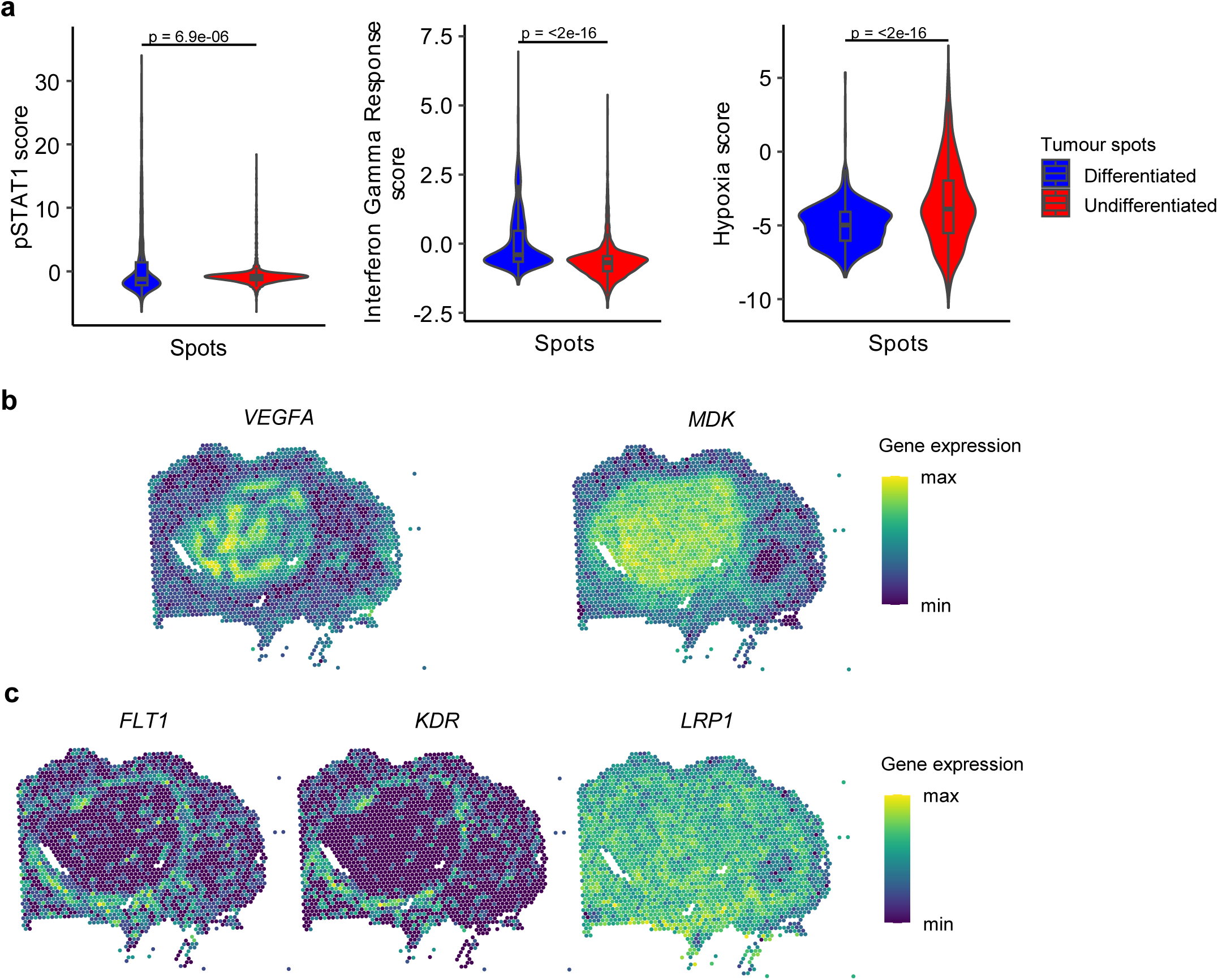
Analyses of environmental signals and selected ligand-receptor pairs. **a** Violin boxplots of the pSTAT1 score, the Interferon Gamma Response hallmark score, and of the hypoxia score in differentiated and undifferentiated tumour spots, in all 4 tumours merged. Mann-Whitney U tests, p-values are indicated on the boxplots. **b** Spatialplots of ligand-coding genes *VEGFA*, and *MDK* (from undifferentiated spots) on patient #1 sample. **c** Spatialplots of receptor-coding genes *FLT1*, *KDR* (receptors of VEGFA), and *LRP1* (receptor of MDK) on patient #1 sample.

**Extended Data Fig.12 (related to Fig.5):**
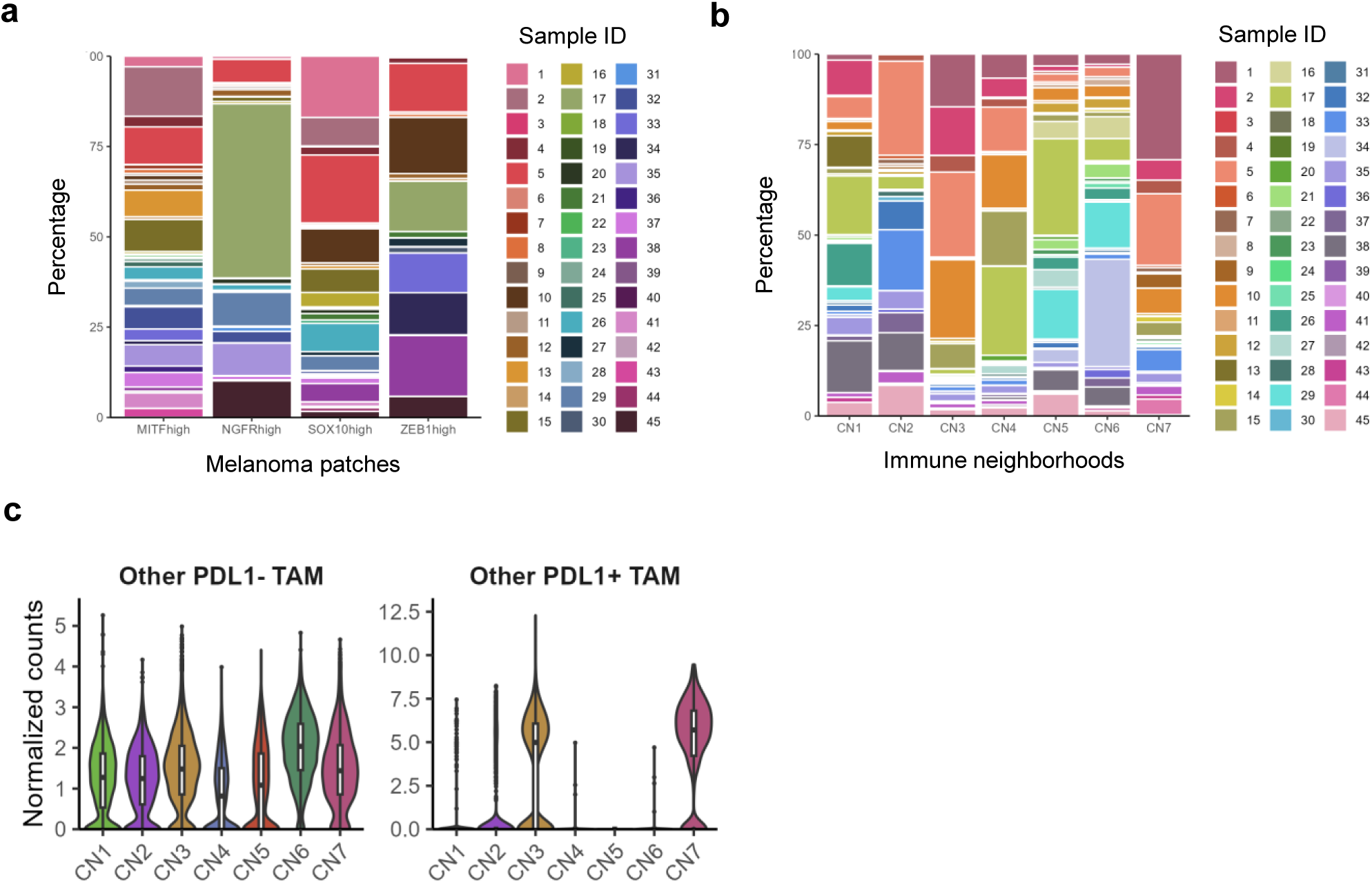
Patient contribution and composition of immune cellular neighbourhoods. **a** Melanoma patches contribution per sample. **b** CN contribution per sample. **c** Violin boxplots of the normalized counts of other TAM across the 7 CNs.

**Extended Data Figure 13 (related to Fig.6):**
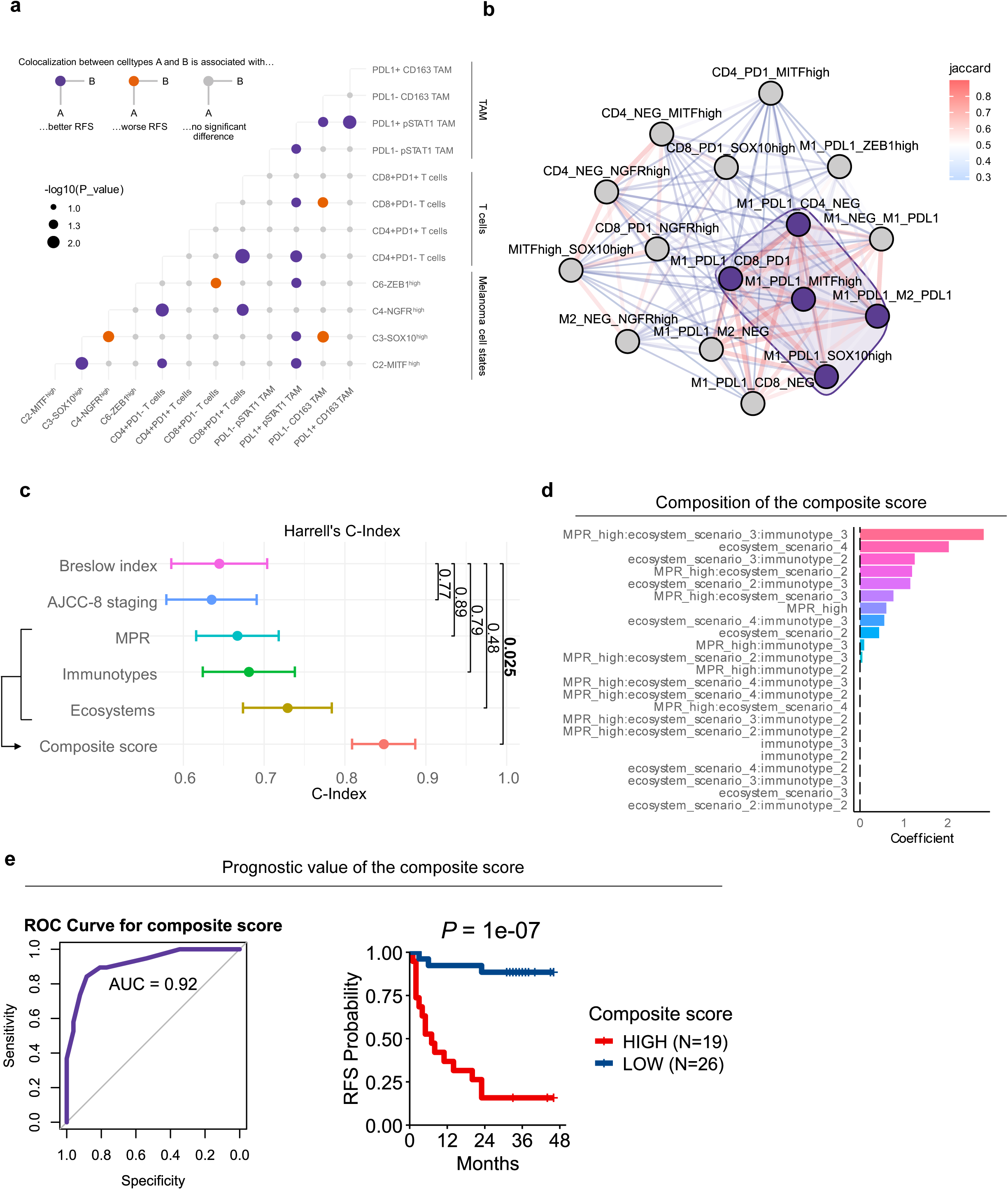
Identification of tumour-immune ecosystems and their prognostic impact in patients. **a** Bubbleplot of the association between pairwise spatial co-localization of TAM, T cells and melanoma subsets with recurrence-free survival (RFS). The points represent the −log10(p-value) of a univariate Cox proportional hazard regressed on the co-localization scores. Each pairwise pattern is either associated with a significantly improved (in purple) or worsened (in orange) RFS. Gray dots are interactions that are not significantly associated with changes in RFS. Every cell types involved in detrimental pairwise patterns (in orange) are then included in the ecosystem U1 in (Fig.6c). **b** Weighted graph of the pairwise cellular co-localizations associated with the relapse-free (RF) status of patients at 2 years. Each node represents a pairwise interaction. The thickness of the edges between two nodes increases with the Jaccard’s similarity index, as well as the colour palette. The nodes are clustered through Leiden clustering based on the Jaccard index to identify highly recurrent pairwise co-localization patterns. A single cluster of 5 pairwise patterns (in purple) is identified, the cell types of which forming the ecosystem D1 in (Fig.6d). M1 represents pSTAT1+ TAM, whereas M2 represents CD163+ TAM. **c** Harell’s C-index of the different metrics as compared to the Breslow index. The composite score is computed from the MPR, immunotypes and ecosystems (see Methods). The C-index with its standard deviation is shown for each metric. P-values are from Mann-Whitney U tests compared to Breslow index. **d** Coefficients of the features included in a Lasso regularized Cox proportional hazard model to build the composite score. Features with a positive coefficient are associated with worsened relapse-free survival (RFS). Features with non-null coefficients are subsequently included in the composite score. **e** Receiver Operator Curve for the composite score regarding the prediction of the 2-year status (left) and Kaplan-Meier curve of the 45 patients separated by the composite score (right). The p-value is from a Log-rank test.

**Extended Data Figure 14:**
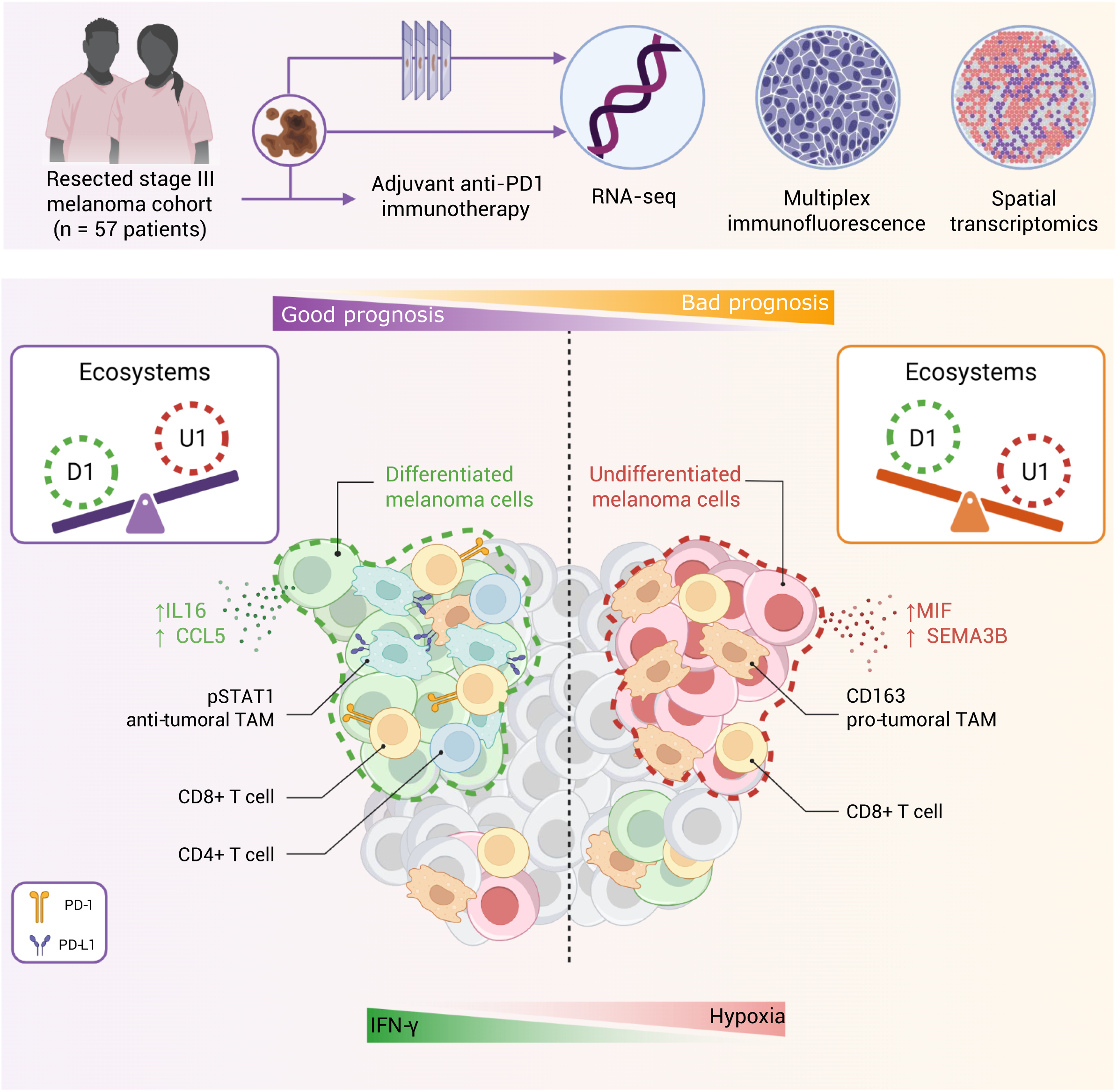
Graphical abstract. Created with BioRender.

**Extended Data Table 1.**
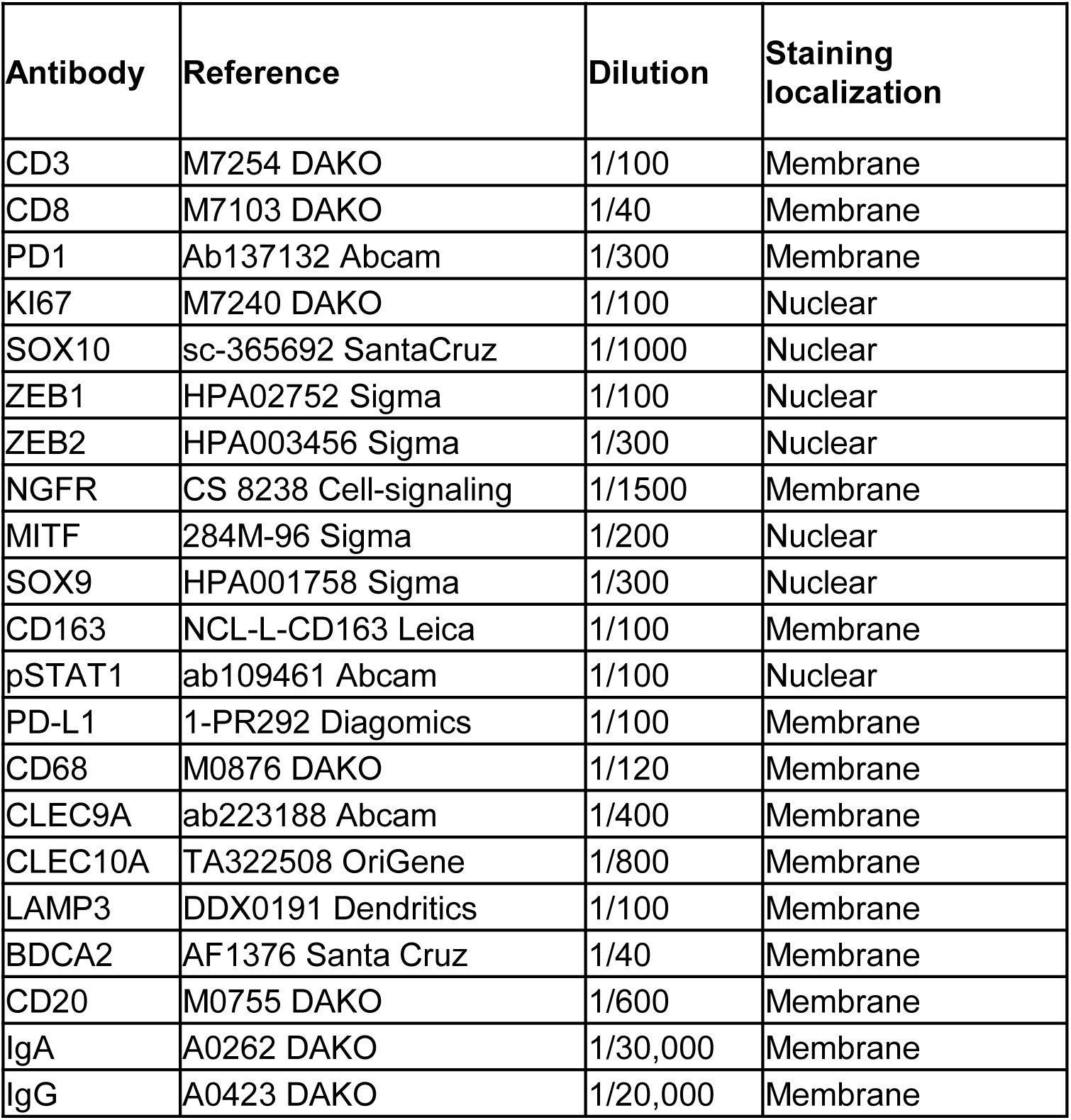
Antibodies used for multiplex immunofluorescence stainings.

**Extended Data Table 2.** pSTAT1 signature.

| Genes |
| --- |
| *IFI44L* |
| *IFI27* |
| *IFI44* |
| *ISG15* |
| *RSAD2* |
| *SIGLEC1* |
| *IFIT1* |
| *ISG15* |
| *GBP1* |
| *IFIT2* |
| *IRF1* |
| *APOL6* |
| *OAS1* |

